# Cardiac interstitial tetraploid cells that escape replicative senescence are found in rodents but not large mammals

**DOI:** 10.1101/521716

**Authors:** Kathleen Broughton, Tiffany Khieu, Nicky Nguyen, Michael Rosa, Sadia Mohsin, Pearl Quijada, Bingyan Wang, Oscar Echeagaray, Dieter Kubli, Taeyong Kim, Fareheh Firouzi, Megan Monsanto, Natalie Gude, Robert M. Adamson, Walter P. Dembitsky, Michael Davis, Mark A. Sussman

**Affiliations:** San Diego State University Heart Institute and the Integrated Regenerative Research Institute 5500 Campanile Drive, San Diego, CA 92182; Temple University, Philadelphia, PA; Division of Cardiology, Sharp Hospital, San Diego, CA; Emory University, Atlanta, GA

## Abstract

Ploidy for cardiomyocytes is well described but remains obscure in cardiac interstitial cells (CICs). Ploidy of c-kit+CICs (cCICs) were assessed using a combination of confocal, karyotypic, and flow cytometric assessments coupled with molecular and bioinformatic analyses. Fundamental differences were found between cultured rodent (rat, mouse) cCICs possessing mononuclear tetraploid (4n) content versus large mammal (human, swine) with mononuclear diploid (2n) content. *In-situ* analysis, confirmed with fresh isolates, revealed diploid content in cCICs from human and a mixture of diploid and tetraploid nuclei in mouse. Molecular assessment of the p53 signaling pathway provides a plausible explanation for escape from replicative senescence in rodent but not human cCICs. Single cell transcriptional profiling reveals distinctions between diploid versus tetraploid populations in mouse cCICs, alluding to functional divergences. Collectively, these data reveal fundamental species-specific biological differences in cCICs that could account for challenges in extrapolation of myocardial preclinical studies from rodent to large animal models.

## Introduction

Ploidy, the number of chromosomes within a cell, is maintained as diploid sets in most mammalian somatic tissues level. Although diploid chromosome content is often considered as ‘normal’, higher levels of ploidy with multiple chromosome sets referred to as “polyploid” are widespread across phyla as well as diverse organisms including mammals. Polyploidy was first observed in plants over a hundred years ago and is an advanced evolutionary trait of adaptation and survival^1,2^, evidenced by polyploid genomic content in over 70% of flowering plants^3^. Polyploidization is often a consequence of normal development, aging, disease, and tissue regeneration processes^4,5^. Polyploid DNA content in mammals normally arises during fetal development in the placenta (trophoblasts)^6^ or in postnatal development of the liver (hepatocytes) ^5,7^ bone marrow (megakaryocytes)^8^, skin (karatinocytes)^9^ and during aging and disease within the liver and heart (cardiomyocytes) ^5,7,10^. Generation of polyploid chromosome content occurs through initiation of abortive mitotic activity leading to increased chromosome content within a single nucleus (endoreplication) or generation of multiple nuclei (endomitosis)^11,12^. Biological significance of such ploidy variations has been investigated for decades spawning substantial speculation, yet the fundamental biological impetus for genome multiplication remains surprisingly obscure.

Decades of research has uncovered many instances where polyploidization is intimately linked to biological processes in living organisms. Polyploidization creates genetic diversity enabling adaptation to environmental stress^13^. Ploidy influences cellular biological properties in multiple ways, and presence of polyploid cells in mammalian organisms indicates evolutionary conservation consistent with a fundamental biological role^5,12,14^. Several beneficial traits have emerged to account for initiation of polyploidization including: adaptation to environmental stress, cell cycle regulation, DNA damage resistance, abrogation of senescence and apoptosis, tissue repair and regeneration^5,7^. Many of these examples occur in plants and invertebrate species where regenerative capabilities are a normal part of their biology (plants regenerate from cuttings, planaria can be cut in half and regenerate^1,2^). Similarly, many lower vertebrates with remarkable regenerative properties possess polyploid DNA content (salamanders, fish, frogs^1,4^) or genome duplication (zebrafish^15^). It may not be entirely coincidental that some of the most regenerative tissues in the adult human body including liver, skeletal muscle, and skin all possess polyploid cellular populations.

Although polyploidy is characteristic of the mammalian myocardium, the adult heart is notoriously refractory to *de novo* cardiomyogenesis that has been correlated to increasing ploidy content^16,17^. Indeed, cellular and nuclear ploidy in adult mammalian cardiomyocytes is in a dynamic state, dependent upon genetics, age and environmental circumstances^10,16,18,19^. Binucleated cardiomyocytes are notoriously refractory to proliferation stimuli, perhaps serving a role in maintaining cell function and/or adaptation to stress^13^. The fluid nature of ploidy in the adult mammalian heart suggests functional involvement, albeit poorly understood, in response to injury and aging. Surprisingly, despite abundant recognition of ploidy as an inherent biological property of the adult mammalian cardiomyocytes, ploidy status of the cardiac nonmyocyte population and biological significance of ploidy in myocardial interstitial cells has received relatively little attention.

The cardiac interstitial cell (CIC) population is a heterogeneous collection of cell types including (but not limited to) fibroblasts, stromal cells, and various progenitor / stem cell populations^20,21^. The CIC population serves a critical role in both homeostasis as well as response to injury although ploidy shifts in CICs remain essentially unstudied. Among CIC cell types, c-kit+ CICs proliferate in response to infarction injury by transient proliferation as reported by our group^22^. The impact of myocardial infarction injury upon CICs, including that of c-kit+ cells, has been extensively studied to exquisite detail for decades as summarized in many research studies and reviews^23–26^. Based upon our extensive experience studying c-kit+ cells for over a decade^21,22,27–30^, we chose to focus our initial investigation upon furthering understanding of ploidy in this subset of the CIC population. Functional consequences of polyploidy in these interstitial cells may correlate with transcriptional changes leading to genetic diversity, adaptation to injury, and improved survival in stress response. Mouse mesenchymal stem cells (mMSCs) can exist as tetraploids and cultured tetraploid mMSCs demonstrate reduced p53 activity compared to diploid mMSCs^31^. In our laboratory, mMSCs fused with mouse cultured ckit+ cardiac interstitial cells, termed cardiac stem cells (CSCs) resulted in chimeric stem cells (CardioChimeras) with sustained enhanced support for functional repair after infarction injury compared to either parent line^32^. In reviewing these studies, we noticed that CardioChimeras with higher chromosome counts correlated with better functional capacity *in vitro* and *in vivo*. The CardioChimera study prompted further investigation of cardiac stem cell ploidy differences between rodents, large mammals and human due to striking differences between species.

Presence of polyploid cells in the interstitial cell population suggests a novel aspect of heterogeneity previously unappreciated and could account for observations of remarkable myocardial reparative potential in murine experimental model systems. Profound interstitial cell expansion and acute remodeling to reinforce ventricular wall integrity after massive infarction injury is a survivable and tolerable event in rodents, whereas similar levels of damage ends in death or rapid adverse remodeling in larger mammals. As polyploid cells are demonstrably correlated to specialized function in tissues and organisms, this study centers upon DNA and ploidy content within a select subpopulation of cardiac nonmyocyte cells. Multiple rodent (mouse, rat), large mammal (feline, swine) and human samples were studied over development, adulthood and after pathologic injury using *in situ, in vitro* and freshly isolated cell preparations. These data provide compelling evidence for the existence of a previously uncharacterized cardiac-specific tetraploid nonmyocyte interstitial cell population in rodents as well as a role for ploidy to influence physiologic response to environmental stress and pathological injury.

## Results

### Frequency of higher ploidy in human cardiomyocytes increases with age and disease

Ploidy is increased in adult human cardiomyocytes isolated from heart failure patients relative to normal control samples^33^. *In situ* ploidy analysis was performed through quantification of diamidino-2-phenylindole (DAPI) fluorescence intensity of reconstructed confocal z-stacks^34^. Ploidy of single nuclei from cCICs and cardiomyocytes were assessed by *in situ* DAPI fluorescence intensity of reconstructed confocal z-stacks from fetal (Figure 1 A), normal adult (Figure 1 B,C) and heart failure patients (Figure 1 D,E). Ploidy grouping were determined by frequency of cells within fluorescence intensity ranges and results are reported in the aggregate from multiple patients and experiments (Figure 1F). Diploid content was predominant in cCICs from fetal, normal adult and LVAD adult tissues. Cardiomyocytes in fetal samples demonstrate a diploid content, whereas normal adult cardiomyocytes are approximately 71% diploid with 29% tetraploid and LVAD adult cardiomyocytes demonstrated the greatest variation in ploidy content with approximately 30% diploid, 56% tetraploid and 14% greater than tetraploid (Figure 1 G,H). These results demonstrate the ckit+ CICs demonstrate less ploidy variation with aging or disease compared to cardiomyocytes in the human heart.

**Figure 1:**
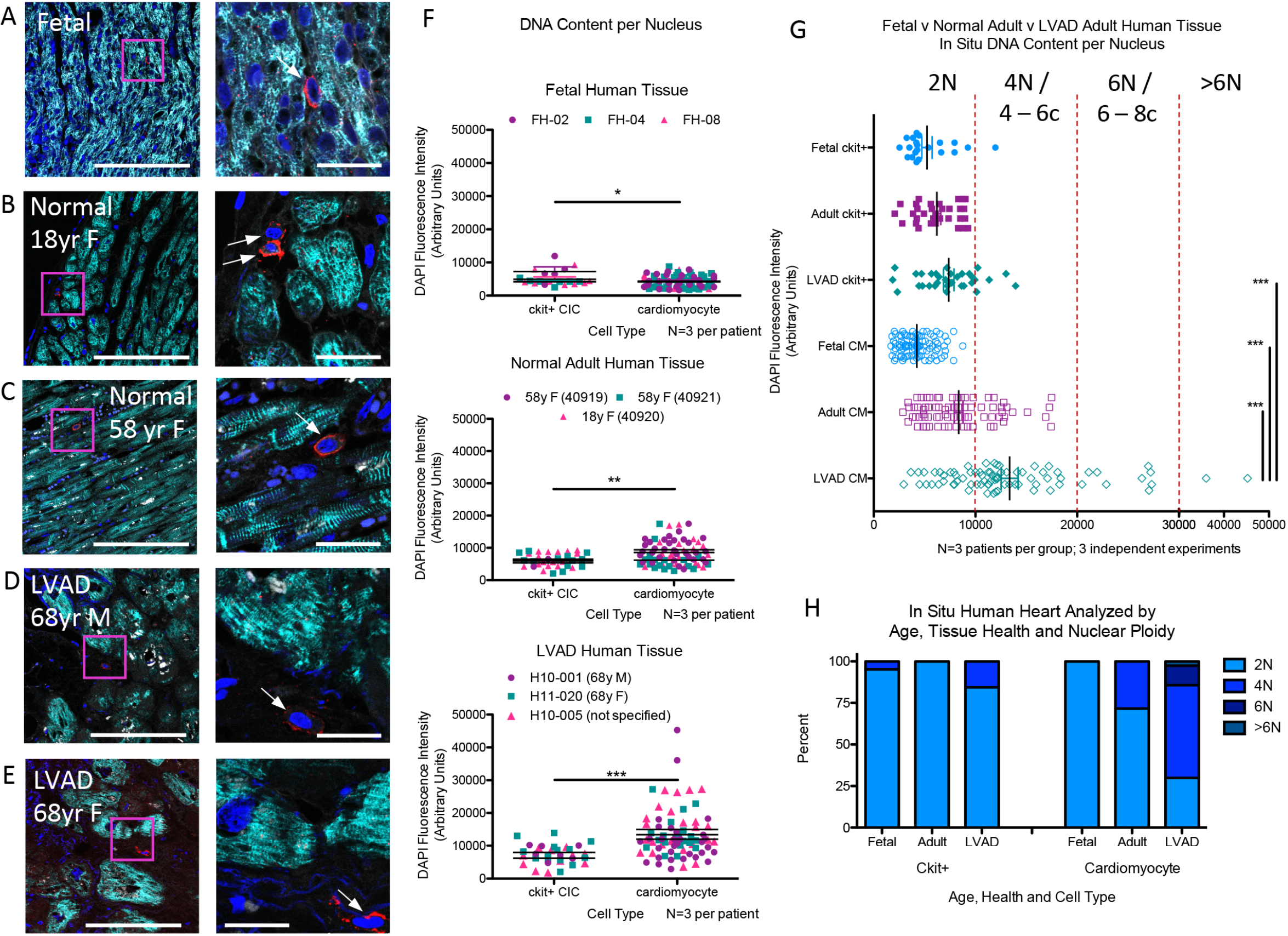
Higher ploidy level increases with age and disease in human cardiomyocytes. Human myocardial tissue sections assessed by *in situ* quantitation of ploidy level from 16 week old fetal (A,), 18 (B) and 58 (C) year old female with normal heart histology and function, and 68 year old male (D) and female (E) explanted from the left ventricle free wall from heart failure patients receiving LVAD implant (zoomed out scalebar = 150 um; zoomed in scalebar = 25 um). Boxed regions of higher magnification are shown to the right of each scan. Arrows point to example cCICs included in the analysis. Quantitation of *in situ* DNA content per nucleus of cardiomyocytes and ckit+ cells measured by DAPI fluorescent intensity of nucleus within 3D reconstruction of tissue, analyzed by t-test (F). Compiled data of *in situ* DNA content per nucleus of cardiomyocytes and ckit+ cells measured by DAPI fluorescent intensity of nucleus within 3D reconstruction of tissue (G). Percent of cCICs and cardiomyocytes nuclei with diploid tetraploid and higher ploidy content (H). *P<0.05; **P<0.01; ***P<0.001. Data are presented as Mean±SEM and analyzed using a t-test (F) or one-way ANOVA with Bonferroni post-hoc test (G).

### Multiple ploidy populations in murine cardiomyocytes and c-kit+ CICs emerge during postnatal stages to adulthood

Ploidy content within mononucleated adult murine cardiomyocytes is predominantly mononuclear diploid, transitioning to binucleation during development^16^. Postnatal proliferation of murine cardiomyocytes and CICs occurs primarily within the first week after birth^35^ as cardiomyocytes acquire variation in ploidy, but ploidy of the CICs remains obscure. Ploidy of cCICs and cardiomyocytes was assessed by *in situ* DAPI fluorescent intensity analysis at postnatal ages of 3 days, 7 days, 1 month and 3 months (Figure 2). cCICs demonstrated approximately 75% diploid content at the 3, and 7 days post birth with 47% and 61% diploid at 30 and 90 days post birth, respectively. Single nuclei of cardiomyocytes display approximately 58% diploid content at 3 days post birth, which became approximately 85% diploid by day 7 postnatal and maintained at that level at 1 and 3 months (Figure 2E,F). Cardiomyocytes undergo DNA doubling and a proliferative burst during the first few days after birth^36^, consistent with increased number of cardiomyocytes possessing tetraploid nuclei at 3 days post birth. These results demonstrate diploid, tetraploid and a small contribution of cells with a higher ploidy within adult cCICs and the ploidy transition of single nuclei in cardiomyocytes between the neonate to adult state during postnatal development.

**Figure 2:**
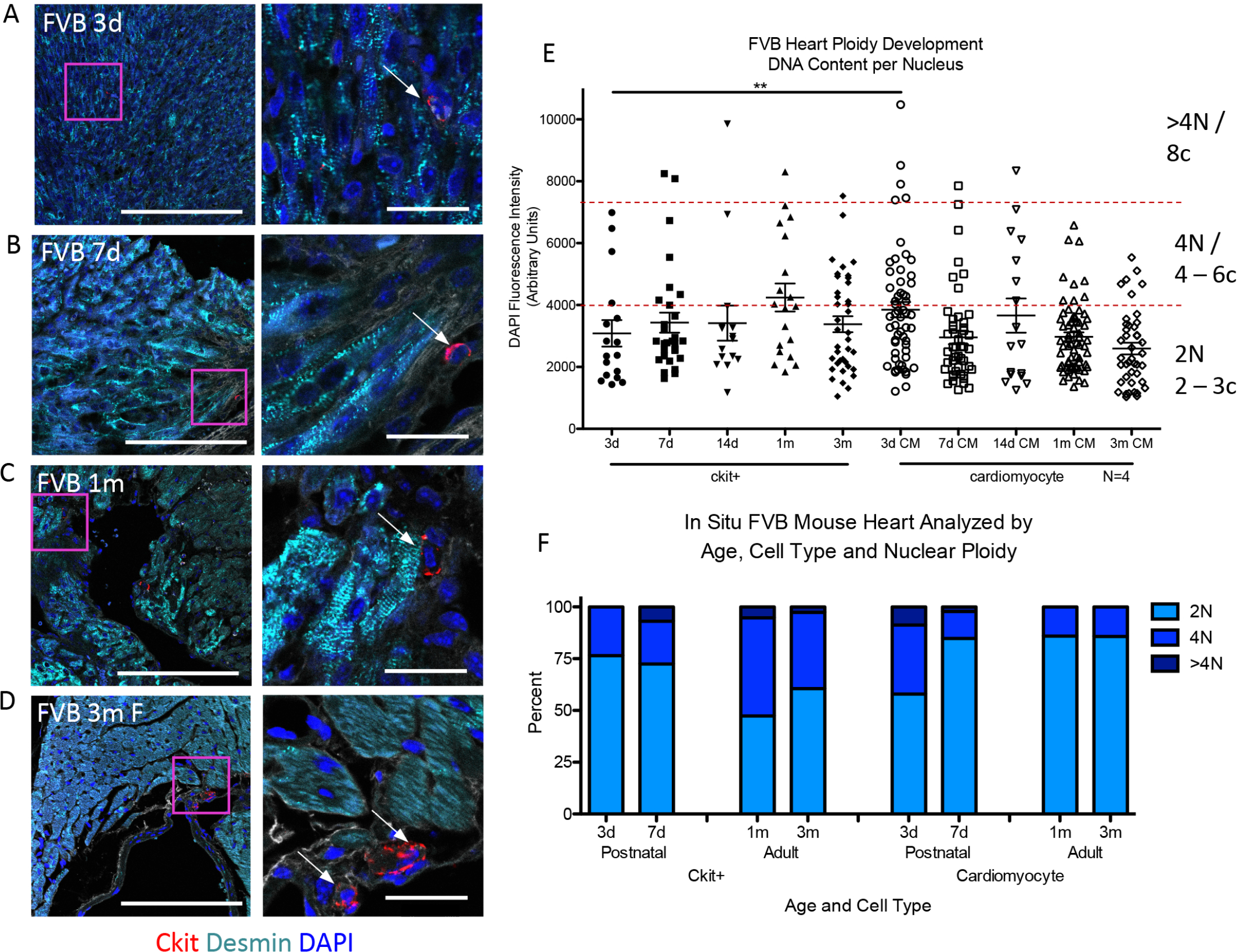
Ploidy variability in murine cardiomyocytes and c-kit+ CICs. FVB mouse myocardial tissue sections assessed by *in situ* quantitation of ploidy level post birth at 3 days (A), 7 days (B), 30 days (C) and 90 days (D) (zoomed out scalebar = 150 um; zoomed in scalebar = 25 um). Boxed regions of higher magnification are shown to the right of each scan. Arrows point to example cCICs included in the analysis. Compiled quantification of *in situ* DNA content per nucleus of cardiomyocytes and ckit+ cells measured by DAPI fluorescent intensity of nucleus within 3D reconstruction of tissue (E). Percent of cCICs and cardiomyocytes nuclei with diploid tetraploid and higher ploidy content (F). **P<0.01. Data are presented as Mean±SEM and analyzed using a t-test, at each time point, or one-way ANOVA with Bonferroni post-hoc test within cell types (E).

### Tetraploid content of endogenous ckit+ CICs uniquely observed the murine heart

Ploidy state of ckit+ cells within multiple adult FVB mouse tissues was determined by *in situ* DNA analysis of heart, intestine and bone marrow. Ploidy of mononuclear cCICs show both diploid and tetraploid cells, whereas diploid content predominates within ckit+ cells of the intestine and bone marrow (Figure 3A-C). C-kit+ CICs and bone marrow stem cells (BMSCs) were isolated, expanded and assessed for ploidy after culturing, as our prior study reveals substantial transcriptional reprogramming following culturing of stem cells^21^ (cultured ckit+ CICs referred to as cardiac stem cells (CSC)). Marked divergence in ploidy levels was evident in CSC cultures relative to BMSCs of adult FVB mice showing tetraploid versus diploid content, respectively, by confocal (Figure 3D,E) and flow cytometry (Figure 3F) analyses. Karyotyping verified CSC mononuclear tetraploid (Figure 4A) versus diploid content of BMSCs and bone marrow-derived MSCs (Figure 3G,H). Collectively, these findings highlight distinct tetraploid content of c-kit+ CIC and CSCs relative to c-kit+ cells from non-cardiac murine tissue sources or cultured bone marrow stem cells.

**Figure 3:**
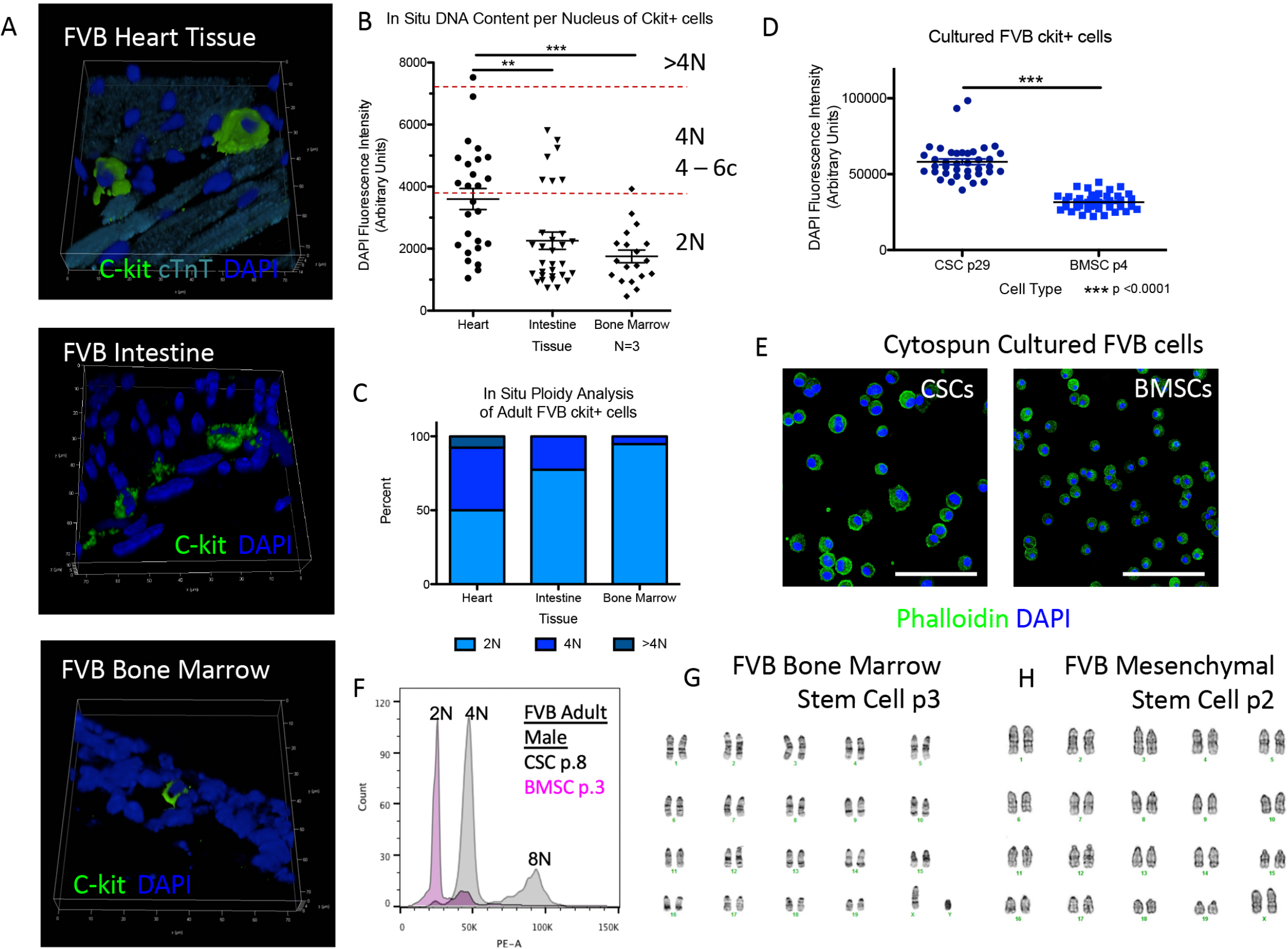
Tetraploidy is characteristic of an endogenous c-kit+ CIC subpopulation. FVB mouse myocardial, intestine and bone marrow tissue sections assessed by *in situ* quantitation of ploidy level at 90 days post birth (A). Compiled quantification of *in situ* DNA content measured by DAPI fluorescent intensity of nucleus within 3D reconstruction of tissue demonstrate cardiac ckit+ CICs have higher ploidy levels compared to ckit+ from intestine or bone marrow (B). Percent of ckit+ cells nuclei with diploid tetraploid and higher ploidy content from myocardial, intestine and bone marrow tissue (C). Ckit+ cells isolated and cultured from cardiac tissue demonstrate higher ploidy levels compared to cultured ckit+ cells from bone marrow, (D), measured by DAPI fluorescent intensity of nucleus within 3D reconstruction of tissue (scalebar = 150 um) (E) and propidium iodine fluorescent intensity of nucleus using flow cytometry (F). G-band karyotype analysis verify diploid content of cultured ckit+ bone marrow stem cells (G) and bone marrow-derived mesenchymal stem cells (H). **P<0.01; ***P<0.001. Data are presented as Mean±SEM and analyzed using a t-test (D) or one-way ANOVA with Bonferroni post-hoc test (B).

**Figure 4:**
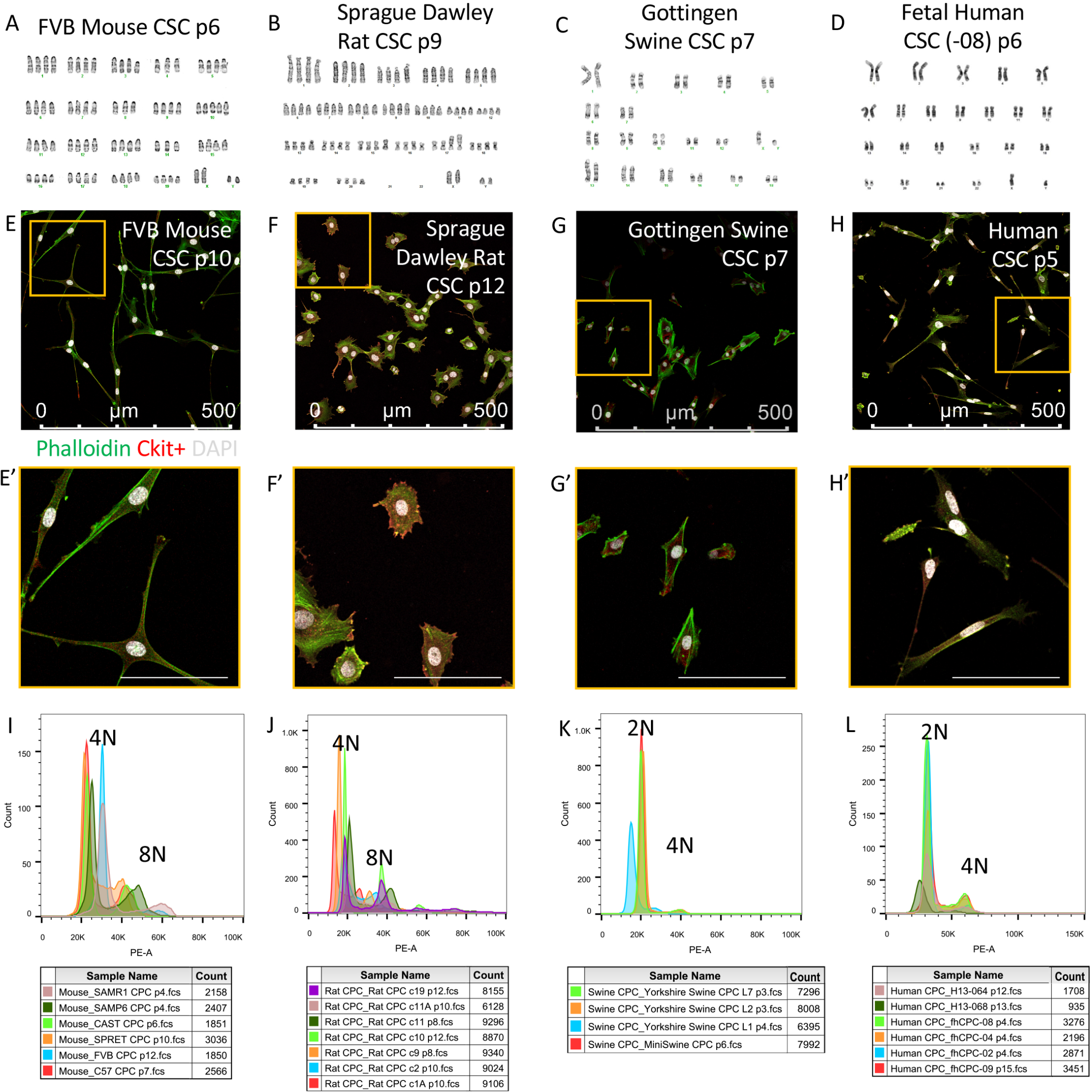
Mononuclear tetraploid content distinguishes rodent cardiac stem cells *in vitro*. G-band karyotype analysis performed upon cultured CSCs reveals tetraploid content of rodent mouse (A) and rat (B) in contrast to diploid content of swine (C) and human (D) samples. Immunocytochemistry of CSC verify mononuclear content of karyotyped cells (E-H) and zoomed in images (scalebar = 100 um) (E’-H’). Multiple sample from different CSC lines verify consistent tetraploid content in adult mouse (I) and rat (J) CSCs, while swine (K) and human (L) CSCs are consistently diploid.

### Rodent CSCs acquire uniformly mononuclear tetraploid content *in vitro*, in contrast to diploid cells of pig or human origin

Additional evidence to support aforementioned ploidy findings was derived from characterization of cell cultures expanded from human, large animal and rodent samples. Six distinct laboratory mouse strains (FVB, C57/B6, CAST, SPRET, SAMP6, SAMR1) were used for CSCs isolation and expansion as well as multiple clonal expanded lines from Sprague Dawley rat samples. Human CSCs were isolated from fetal, pediatric and LVAD patient cardiac tissue. Large animal CSCs were represented by Gottingen and Yorkshire swine as well as domestic feline samples. Karyotype results demonstrate mononuclear tetraploid content in rodent CSCs (Figure 4A,B and S1A,B), whereas human, swine and feline CSCs are mononuclear diploid (Figure 4 C,D and S1C-F). Mononuclear content per cell was verified by immunocytochemistry (Figure 4E-H, S2A-O). Furthermore, karyotype findings for ploidy of each species were consistent with flow cytometry analyses (Figure 4I-L). These results reinforce the finding of distinct mononuclear tetraploidy in rodent CSCs versus the typical diploid content found in human and large mammalian cells, both *in situ* and *in vitro*.

### CSCs ploidy correlates with proliferative capacity

Morphologic and proliferative characteristics of CSCs were assessed *in vitro*. Cell surface area and major to minor axis ratio are similar within each species (Figure S2P-S). Likewise, proliferation rates for each species assayed were comparable (Figure S2T-W). Replicative potential between human and murine CSCs was assessed with continuous passaging until senescence-mediated arrest. Proliferation ceases between passages 12-26 for fetal human CSCs and between passages 9-15 for adult human CSCs isolated from heart failure patients. In contrast, replicative senescence was never achieved with mouse or rat CSCs, with proliferative capacity extending beyond 50 passages. In comparison, mouse BMSCs cease proliferation between passage 4-6. Average growth rate and doubling time for human and standard lab mouse strain (FVB, C57) CSCs demonstrate similar profiles for the first three days after plating with differences only found at day four (Figure S3A,B). Morphology and ploidy of FVB CSCs remained stable between early and late passages (Figure S4A-D) while proliferation rate increased with early long term passaging and then leveling off between passage 50 and 100 (Figure S4E,F). Cell cycle related genes analyzed using RT-PCR reveal significant upregulation of Ccna2, Ccnb2, and Cdc25c in murine CSCs comparing passages 11 versus 100 (Figure S4G). Increased Cdc25c expression in high passage CSC was confirmed by immunoblot (Figure S4H). Murine CSCs underwent cell cycle arrest and lineage commitment towards smooth muscle, endothelial and early cardiac phenotype at both low and high passages after treatment with dexamethasone (data not shown). Collectively, these results establish differences in proliferative capacity with tetraploid CSCs escaping typical replicative senescence arrest normally observed in diploid rodent BMSCs and human CSCs.

### Human and mouse CSCs express senescence-associated p53-associated markers

Tetraploid cells ordinarily senesce in response to endoreplication error^37^, but tetraploid mCSCs have escaped this checkpoint. Cellular senescence markers include p53 (Trp53), p16 (Cdkn2a), and p21 (Cdkn1a)^38^. P53 is associated with the inhibition of cellular growth and subsequent apoptosis; mdm2 controls p53 cell-cycle arrest through mdm2-p53 interaction^39^. To understand the continued growth of tetraploid mCSCs but senescence of hCSCs, a gene panel array of senescence markers and protein was assessed in hCSCs and mCSCs at low versus high passage points (Figure S5A,B). Expression of p53 pathway genes were analyzed by RT-PCR and proteins were measured by immunoblot. Genes upregulated in the high passage compared to low passage hCSCs include apoptosis marker BCL2 and endothelial growth factor receptor KDR (Figure S5A), while genes upregulated in the high passage compared to low passage mCSCs include Cdk2 to regulate S-G2 cell cycle and anti-apoptosis marker Hspb1 while Cdkn1a, Cdkn2a, Mdm2 are downregulated (Figure S5B). In terms of protein, high passage hCSCs express increased total p53 and p53 phosphorylated at Serine 15 (Figure S5C), known to activate p53 [ref], while MDM2 remains constant at low and high passages (Figure S5D). In mCSCs, protein expression of total p53, phosphorylated p53 at Serine 15 and MDM2 is decreased in high versus low passage (Figure S5E,F). Collectively, these data provide insights into the cell-cycle and p53-associated senescence regulation of murine CSCs that evidently override p53-mediated cell cycle arrest.

### Murine CSCs increase negative p53 feedback loop with increased passaging

Tripartite motif-containing (TRIM) superfamily proteins regulate pathogen-recognition and transcriptional pathway defense^40^ by p53 potentiation (TRIM 13,19) or inhibition (TRIM 25,28,29) via the Mdm2-p300-p53 complex^41,42^. TRIM proteins are known to regulate multiple cellular processes including proliferation, differentiation, development, autophagy and apoptosis in mammalian cells^43,44^. Analysis of TRIM expression in hCSCs at replicative senescence revealed increased TRIM 13 transcription together with decreased TRIM 25 and 28 transcription and protein expression (Figure S6A,C,D). Conversely, high passage mCSCs show transcriptional inhibition for TRIM 13 and 19 with concomitant increases in TRIM 25, 28 and 29 (Figure S6B). Protein expression for each TRIM family member was down regulated in high passage mCSCs (Figure S6E,F) and may result as high passage mCSCs demonstrate decreased total p53 and p53 phosphorylated at Serine 15 (Figure S5B). TRIM 29 transcription increased in both human and murine CSCs at higher passage and further studies would be necessary to determine the significance of this finding. These data provide additional evidence that p53 is present in both hCSCs as well as mCSCs at higher passage points but could be subject to TRIM-mediated transcriptional influences upon activity level.

### Freshly isolated murine cardiac lin- c-kit+ CIC exhibit ploidy variation

Ploidy analysis of freshly isolated hematopoietic lineage (lin)- ckit+ viable CICs was performed using Vybrant dyecycle green sorted by flow cytometry. Tetraploid CSCs, verified with karyotype (Figure 4A), were used as a 4N gating control (Figure 5A) for determination of appropriate gates for freshly isolated, viable, FVB lin-ckit+ ploidy populations. Freshly isolated cCIC exhibit two primary ploidy populations (Figure 5B) compared to pure tetraploid cultured mCSC (Figure 5C, overlay). Viable, freshly isolated cCIC are a mixture of diploid (54.51±3.47%), tetraploid (41.8±3.04%), and higher ploidy levels (3.69±0.56%) (Figure 5D; 22000±2915 cells analyzed per experiment). Collectively, ploidy variation findings are comparable for *in situ* and fresh isolation determinations whereas cultured murine c-kit+ CIC are uniformly tetraploid (Figure 5E), confirming ploidy variation in the endogenous murine c-kit+ CIC and culture conversion of these c-kit CIC into tetraploidy.

### Population subsets of diploid versus tetraploid murine fresh isolate lin- c-kit+ CICs revealed by single cell transcriptional profiling

Single cell RNA sequencing (scRNA-Seq) reveals transcriptome differences within freshly isolated CICs^20^ as well as between fresh isolated cCICs versus cultured CSCs^21^. Although subtle, distinctions between diploid versus tetraploid fresh isolates of cCICs were revealed through scRNA-Seq transcriptional profiling. Multidimensional reduction processing coupled with t-Distributed Stochastic Neighbor Embedding (t-SNE) representation through Seurat R package and Barnes-Hut tree-based algorithm reveals three primary aggregates consistent with: 1) interstitial/fibroblast, 2) endothelial, and 3) lymphocyte lineage (Figure 6A–C). Interestingly, these three subgroups were differentially distributed in diploid and tetraploid fresh isolates for fibroblast (69% versus 38%), endothelial (29% versus 59%), and lymphocyte (2% versus 3%) (Figure 6D).

**Figure 5:**
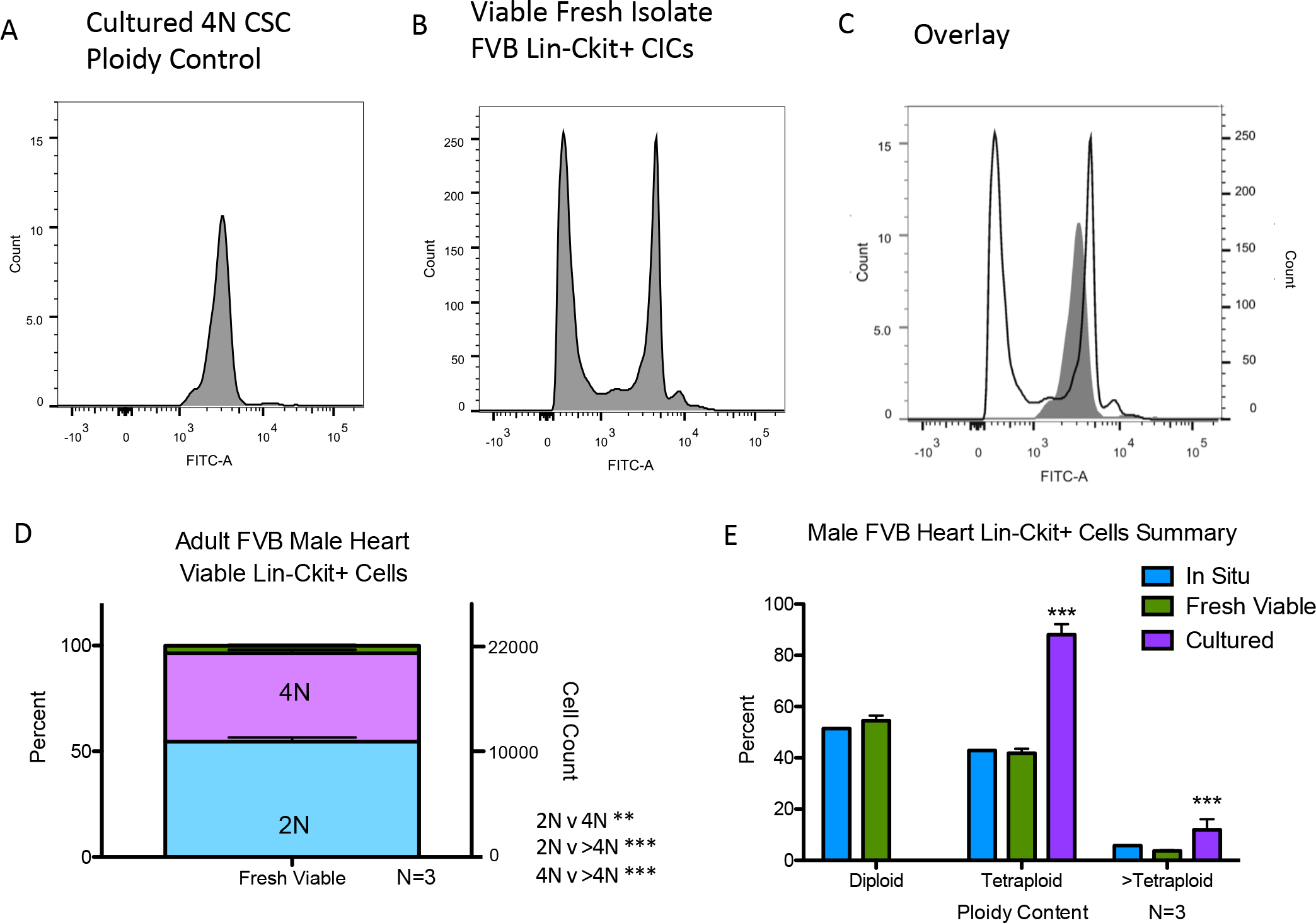
Polyploid states are present in freshly isolated murine c-kit+ CICs. Hematopoietic lineage negative, c-kit positive CICs were freshly isolated and sorted for viability and ploidy content from FVB adult male mice. Using the cultured FVB CSC as a tetraploid control (A), fresh isolates of Lin- ckit+ CICs exhibit a mixture of diploid and tetraploid cells (B) shown by overlay of tetraploid control versus fresh isolate sample (C). Lin-ckit+ CICs possess a similar percentage and cell count distribution of diploid and tetraploid cells with a small fraction of cells with ploidy greater than tetraploid, statistically analyzed using one-way anova (D). Summary of polyploid state from FVB mouse lin-ckit+ cells of the heart, with *in situ* and freshly isolated cells revealing a mixture of mononuclear diploid and tetraploid levels, while cultured CSCs are tetraploid; all groups demonstrated a small fraction of tetraploid cells. **P<0.01; ***P<0.001. Data are presented as Mean±SEM and analyzed using one-way (D) or two-way (E) ANOVA with Bonferroni post-hoc test.

**Figure 6:**
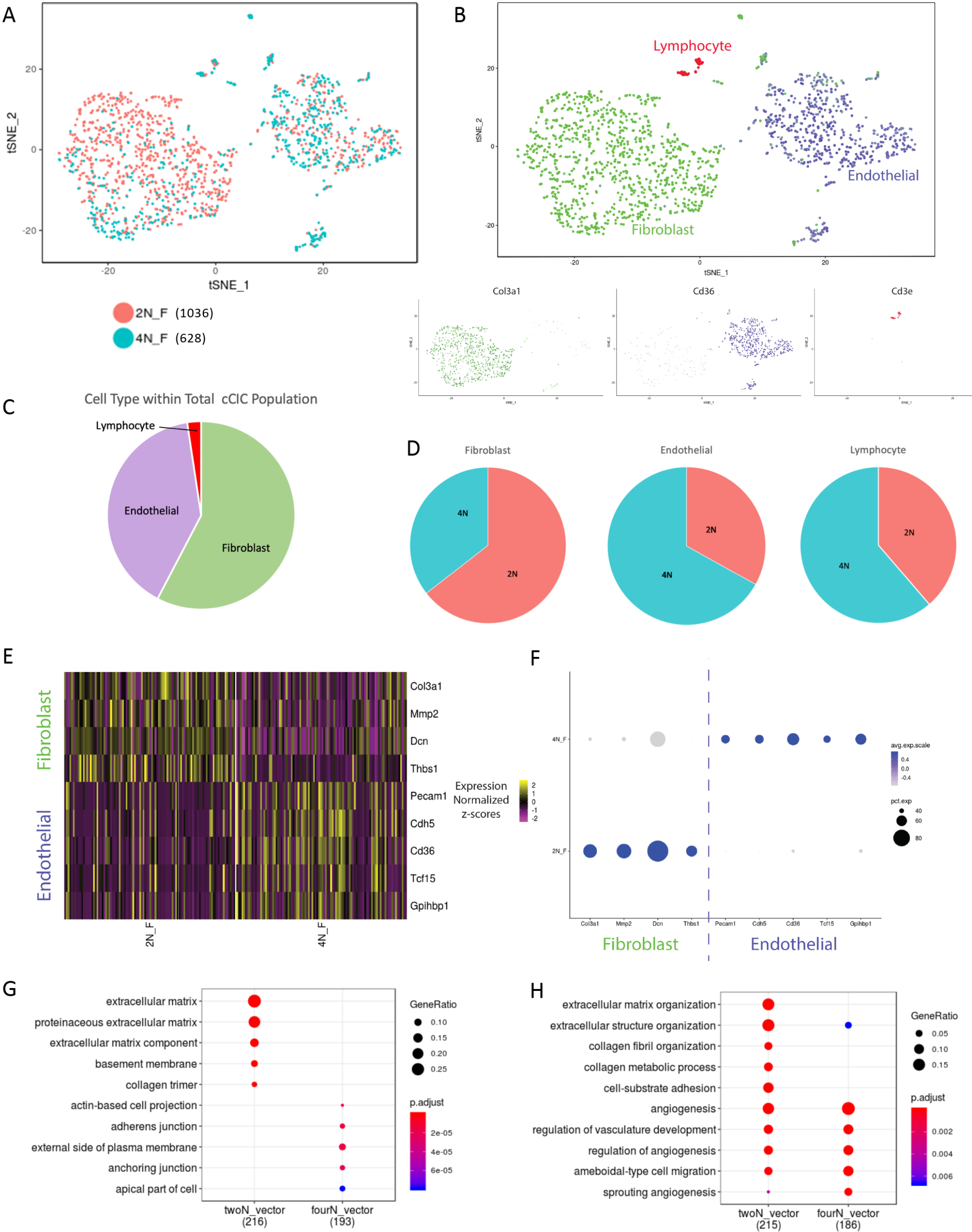
Distinct population characteristics of diploid versus tetraploid fresh murine Lin- c-kit+ isolates revealed by single cell RNA sequencing (scRNA-Seq). Freshly isolated, viable diploid and tetraploid Lin-ckit+ CICs from adult FVB mouse were analyzed using scRNA-Seq. The diploid population (salmon) predominately cluster together, while tetraploid cCICs (teal) cluster in a different cell group (A; cells analyzed per group identified next to ploidy state). Identification of the cell populations demonstrate cCICs are a heterogeneous population of fibroblast, endothelial and lymphocyte cells (B). Percent of cCICs based on cell type (C). Percent of each cell type based on ploidy content (D). Heatmap of upregulated differentially expressed genes (DEGs) specific to endothelial and fibroblast markers confirm transcriptional differences in murine derived diploid (2N_F) and tetraploid populations (4N_F) (E). These DEGs are also displayed in frequency and expression level between the diploid and tetraploid cCICs (F). The top ten gene ontology (GO) terms upregulated in the 2N population display extracellular matrix cellular components while the 4N population cellular component is junction oriented (G). The top ten GO terms by biological process upregulated in the 2N population represents both extracellular matrix and angiogenesis processes, while the 4N population is primarily angiogenesis oriented (H).

A number of basic cellular genes are shared in both the diploid and tetraploid populations; however, 223 differentially expressed genes (DEGs) were identified in the diploid population and 187 DEGs were found in the tetraploid population, with the top fibroblast and endothelial DEGs of each population shown (Figure 6E,F and Supplement Table 1). Extracellular matrix (ECM) and fibroblast gene expression frequency was prominent in the normalized diploid population (Figure S7A,B), identified with Col3a1 (65.72% 2N vs. 34.28% 4N); Mmp2 (65.26% 2N vs. 34.74% 4N); Dcn (55.22% 2N vs. 44.78% 4N); and Thbs1 (69.27% 2N vs. 30.73% 4N). Fibroblast gene expression per cell, shown with violin plots, was elevated in the diploid population (Figure S7C). Endothelial gene expression frequency was higher in the normalized tetraploid population (Figure S8A,B), identified with Pecam1 (34.23% 2N vs. 65.77% 4N); Cdh5 (33.73% 2N vs. 66.27% 4N); Cd36 (35.26% 2N vs. 64.74% 4N); Tcf15 (44.82% 2N vs. 55.18% 4N); and Gpihbp1 (37.5% 2N vs. 62.5% 4N). Violin plots of each gene demonstrates higher endothelial expression per cell within the tetraploid population (Figure S8C). Analysis of the gene onthology (GO) by cellular component verified the 2N population is uniquely linked to ECM-related components (Figure 6G) and biological processes (Figure 6H) compared to the 4N population. These data collectively indicate the majority of 2N lin-ckit+ freshly isolated CICs from murine hearts are fibroblast related to support the extracellular matrix while the majority of fresh isolate 4N lin-ckit+ CICs from murine hearts are endothelial related to support angiogenesis.

### Ploidy reduction occurs in murine lin- c-kit+ CIC following infarction

The cCIC population expands in response to infarction injury^22,45^. Ploidy changes within the cCIC population in response to myocardial infarction was assessed *in situ* in the infarction zone using immunohistochemistry, with non-infarcted hearts as control samples. Mast cells were only identified in infarcted hearts using the surface marker tryptase and were excluded from analysis (Figure 7A,B). Ploidy of cCIC within the infarction and border zone region was identified as over 80% diploid at 4 and 7 days after myocardial infarction with 64% diploid at day 14 post-MI and 92% by day 21 post-MI (Figure 7C) compared to control. These results demonstrate that ploidy of the ckit+ CICs in the infarction zone are predominantly diploid, likely trending toward fibroblast phenotype.

**Figure 7:**
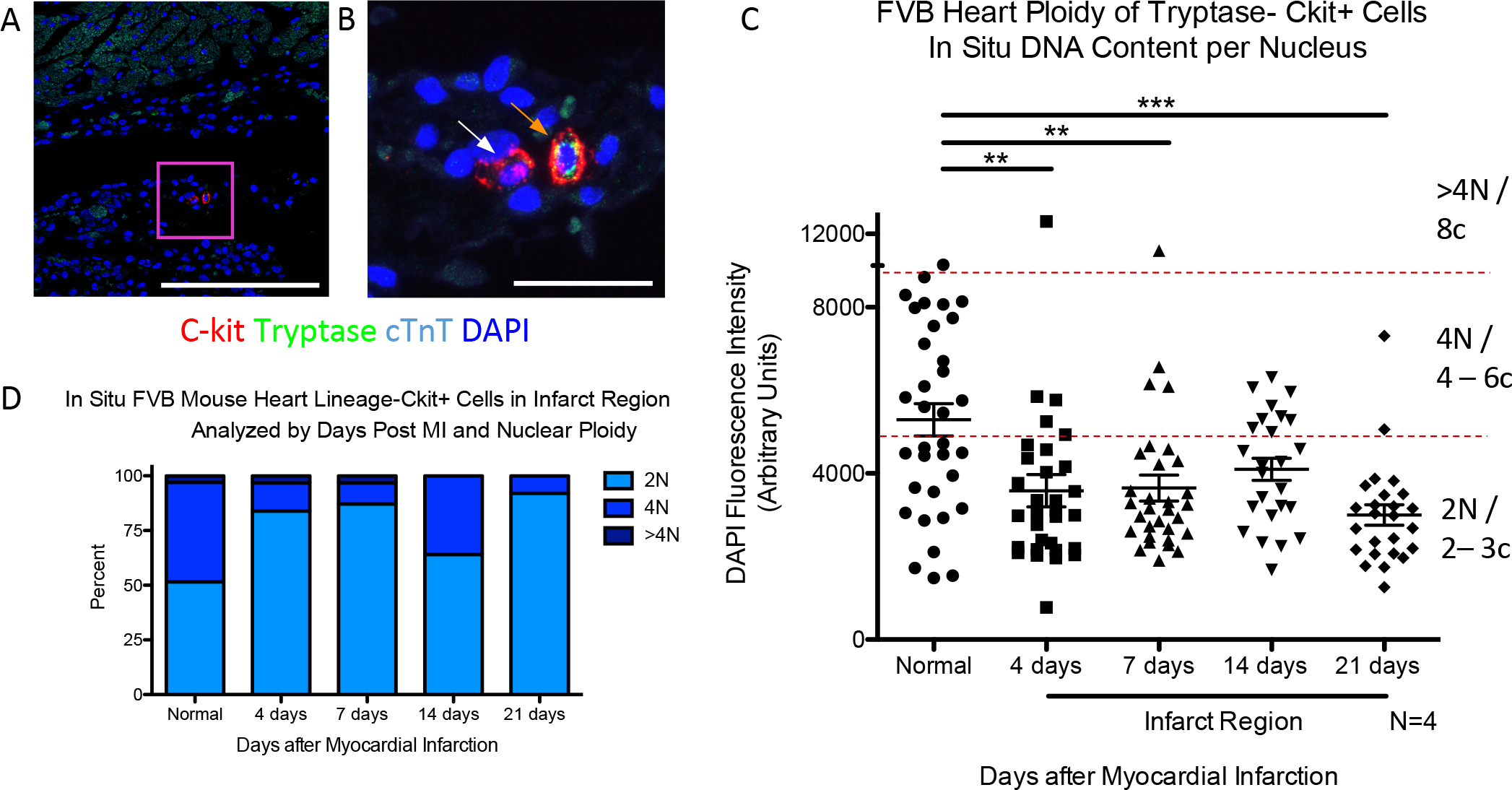
Reduction of DNA content in Lin- c-kit+ CICs following myocardial infarction. Quantitation of *in situ* nuclear DNA content in adult FVB mice following myocardial infarction measured by DAPI fluorescent intensity of tryptase-, ckit+ CIC nuclei. 3D reconstruction of cardiac tissue was performed at 4-, 7-, 14-, and 21-days post injury in the infarction zone. Normal, age, gender and strain matched hearts were used as a control. Tryptase+, a marker of mast cells, were only found in injured hearts within the first week after injury and were not included in the analysis, as shown with tryptase- ckit+ and tryptase+ ckit+ CICs in the infarction zone of an adult, 7-day post-MI FVB heart (A, (scalebar = 150um)) with boxed region of higher magnification (B, scalebar = 25um)). White arrow identifies a cCICs included in the analysis and orange arrow identifies a mast cell, not included in the study. Compiled dot plot of *in situ* DNA content per nucleus of lin-ckit+ cells demonstrates higher frequency of diploid levels in the infarction region at all time points post-MI (C), and is verified upon quantification of percent diploid, tetraploid and greater than tetraploid content (D). **P<0.01; ***P<0.001. Data are presented as Mean±SEM and analyzed using one-way ANOVA with Bonferroni post-hoc test (C).

## Discussion

Polyploidy has been proposed as an advanced evolutionary trait that facilitates regeneration^1^ and provides the capability to cope with stress through possession of extra DNA copies or alternate sets of genes. Indeed, polyploid cells are present throughout human lifespan, with some experts espousing the concept that acquisition of polyploid DNA content is an adaptive process to avoid cell death as well as oncogenic transformation^46,47^. Regenerative activity and functional specialization of cells in mammals are associated with polyploid cell populations^48,49^. In addition, ploidy levels can be influenced as a consequence of aging, stress, acute injury, and environmental conditions^1,2,11,13^. Thus, it is clear that ploidy exists in a dynamic and regulatable state that is consistent with and essential for normal biological function^12,13^. Ploidy variation in cardiomyocytes has been recognized and studied for decades, through binucleation or mononuclear endoreplication, associated with postnatal development, oxidative damage, and telomere erosion^50^. Increased metabolic activity through increased protein production could also be a driver for cardiomyocyte polyploidy, as previously demonstrated in smooth-muscle cells and megakaryocytes^51,52^. To our knowledge, previous ploidy studies of myocardial biology have focused upon cardiomyocytes^5,10,16–19,53^, whereas assessment of CIC ploidy has not been previously addressed, leading to this report that provides several important insights regarding cCIC ploidy content.

A straightforward conclusion from our findings is the clear species and tissue-specific correlation of the tetraploid cCICs in the murine myocardium. Starting with ploidy analysis of *in situ* cardiomyocytes and cCICs performed in rodent (mouse, rat), large mammal (feline, swine) and human samples, the association of tetraploid DNA content with the murine myocardial samples was consistently demonstrable from *in situ*, *in vitro*, and fresh isolation experiments (Figures 2–5). Ploidy state of human cardiomyocyte is a complex, variable, and fluid condition requiring further detailed analyses of more patients and more sample points, similar to conclusions of previously published literature. Previous ploidy studies in the heart are limited to cardiomyocytes. A number of reports have indicated human cardiomyocytes are over 40% mononuclear polyploid by adulthood^54^ and up to 80% polyploid in patients with infarction injury^55^. Results from this study confirm similar findings in the cardiomyocyte population (Figure 1). In murine cardiomyocytes, previous studies have reported that rodent binucleation increases dramatically between postnatal day 4 and 12^56^ and by adulthood, over 80% of single cardiomyocyte nuclei are diploid^57^. Recently, our group reported that postnatal day 2 cardiomyocytes demonstrate the highest number of cardiomyocytes in S/G2/M cell cycle phase during postnatal development^58^. Our study indicates approximately 40% of day 3 single nucleus DNA content of cardiomyocytes are polyploid, which sharply reduces to 15% by day 7 and maintained at 15% into adulthood (Figure 2). This study provides complementary findings to the previous myocyte studies although limitations of the *in situ* study include a general population ploidy analysis based on a limited number of cells, reliance of consistent ploidy content throughout the heart although adult human sections were all from the left ventricle free wall and random selection of cells throughout the heart within the mouse samples. Further studies are needed to understand differences between cellular and nuclear ploidy, as these studies were limited to nuclear content, and further analysis of regional ploidy to determine if ploidy differences exist in each of the four chambers. Future studies will need to incorporate discrimination of ploidy status for individual nuciei within multinucleated or mononuclear cells. Our analysis focused upon determinations for DNA quantitation within individual nuclei rather than nucleation state that together determine cellular ploidy. This issue of cardiomyocyte cell cycle state and ploidy is the focus of ongoing studies and we look forward to exploring these ideas in future investigations.

The observation of tetraploidy in the rodent CSC context was further extended and validated by independent characterization of cultured rat CSCs from the Davis laboratory (Figure 4). Human cCICs are a diploid population from development through adulthood and in pathological states. In comparison, rodent cCICs demonstrate a heterogeneous mononuclear diploid and tetraploid population during development into adulthood. Therefore, profound differences in DNA content exist between cCICs of human and large mammal versus rodent origin, with the reasonable possibility that this fundamental biological difference between species could account, at least in part, for the challenge of extrapolating findings from murine preclinical studies to larger mammals including humans.

Infarction injury in the murine heart favors expansion of the diploid cCIC population in the infarct/border zone. Diploid cCICs predominance may arise from proliferation of diploid cells, ploidy reduction from tetraploid to diploid or a combination of these events. In the Drosophila intestine, polyploid intestinal stem cells (ISCs) undergo amitosis to regenerate new diploid ISCs in response to environmental stress.^59^ Likewise, hepatocytes undergo ploidy reduction during liver regeneration after hepatectomy^60^ In the myocardium, fibroblasts are activated, proliferate and infiltrate in response to myocardial infarction^61^ Considering the majority of diploid cCICs display a fibroblast transcriptome profile, it is likely that cCICs in the infarction are fibroblast. Ongoing studies to delineate CIC ploidy changes in response to infarction are underway.

Another important distinction emerged from *in vitro* rodent ckit+ CICs studies that revealed culture adaptation leads to adoption of a uniform tetraploid phenotype of CSCs capable of escaping replicative senescence, whereas diploid CSCs disappear from the amplified population. Multiple concurrent processes could explain to dominance of the tetraploid population in vitro. Examples include 1) the diploid population undergoing endoreplication or endomitosis to become tetraploid, 2) indirect influence of tetraploids to inhibit diploid proliferation through cell-cell communication, possibly via secreted factors, or 3) greater adaptive capacity of the tetraploid population to the stress of in vitro expansion, leading to diminished competitive capability of the diploids. Indeed, tetraploid cCICs and CSCs may inhibit diploid proliferation through cell-cell communication, as was recently discovered between tetraploid and diploid hepatocytes to maintain stability and prevent tumor formation^62,63^. Although we cannot exclude fusion as a possible mechanism^64^, no evidence of fusion has been observed in time-lapse recordings of CSC cultures (unpublished observation). A future direction of this project is to understand the ploidy shift of mCSC and determine the relationship between *in vitro* and the *in vivo* system.

Functional *in vitro* differences were consistent regardless of species, between diploid versus tetraploid CSCs. Human CSCs from heart failure patients undergo replicative senescence at passage 12-16, whereas pediatric and fetal CSCs typically expand through 21-26 passages. In comparison, rodent CSCs exhibit proliferation without succumbing to replicative senescence. Respective ploidy states of human and rodent CSCs remained invariant throughout passaging showing stability of the divergent ploidy states between the two species. Prior *in vitro* studies involving human and mouse embryonic stem cells (ESCs) demonstrate polyploidy resulting from failed mitosis^65^ In this study, mouse ckit+ BMSCs maintain diploid content after culturing while mCSC demonstrate a pure tetraploid content upon culturing after an initial isolation displaying a mixture of diploid and tetraploid content. Understanding the in vitro ploidy shift in the mCSC from an initial mix of diploid to pure tetraploid population may provide mechanistic insights into replicative senescence shown in hCSCs and mBMSCs.

On a molecular level, rodent CSCs show a profile consistent with inactivation of cell cycle checkpoints together with downregulation of p53 that could be functionally inhibited by differential expression of TRIM family members. The relationship of p53 to suppression of tetraploidy has been reported in murine embryonic development, with p53 downregulation rescuing tetraploid embryos^66^ and providing chromosomal stability for tetraploid cells^67^. Also, p53 plays a role in ploidy resolution during liver regeneration^68^. Furthermore, assessment of DNA content in mesenchymal stem cells derived from p53 knockout mice cultured for 40 days showed conversion to tetraploidy in contrast to normal control cells that remained diploid^69^. It is tempting to speculate that tetraploid CSCs represent an evolutionary adaptation of CICs providing a functional benefit to rodents. The capacity of CICs to avoid cellular senescence without oncogenic transformation would provide a reservoir of cells with enhanced proliferative potential as well as resistance to environmental stress and preservation of functional responsiveness. The down regulation of p53 in high passage mCSCs is coupled with transcriptional upregulation of negative feedback regulation of p53 with TRIM 25, 28, and 29. Protein for these TRIM isoforms are down in high passage mCSCs, which could result as p53 protein is also down in high passage mCSCs. We recognize that mRNA transcript levels do not necessarily correlate with protein expression^70^, as was the case here with select TRIM isoforms for the human and mouse CSCs. Additional studies revealing how various TRIM isoforms respond to environmental stress are clearly warranted to delineate functional, molecular, and mechanistic differences between diploid versus tetraploid CSC to reveal additional insights regarding the capacity of a rodent heart to withstand pathological damage and how larger animals may be not as well equipped on a cellular level.

CIC number increases through postnatal proliferation as well as post-infarction in rodents^35,55,71^ and humans^55^. Early studies such as these assess the CIC population as a single cell type, but the recent advent of scRNA-Seq technology has helped describe heterogeneity of the CIC population normally present in the adult murine heart^17,20^. Findings reported on the transcriptome level point to not only cellular heterogeneity, but a blend of subtle transcriptional profiles within each cell subtype that reflects population diversity^20,21^. In comparing the fresh isolate 2n and 4n hematopoietic lineage negative, ckit positive CICs, results demonstrate differences in the heterogeneity of the two populations. The 2n and 4n freshly isolated lineage-ckit+ CICs from adult murine hearts exhibit differential DEG expression, identifying the 2N cells predominately as fibroblast and ECM-support cells, while the 4N population is endothelial and angiogenesis-support cells (Figure 6). Interesting, lineage-ckit+ CICs in the infarction zone of an adult mouse heart are nearly all diploid, supporting the remodeling of the heart and scar formation with some vascularization (Figure 7).

Ploidy changes is postulated as a rapid and potentially reversible response to environmental stress^72–74^. Multiple explanations have been offered for the biological significance of polyploidy, including the intriguing idea that the polyploidy appears to be permissive for recovery from the senescence-arrested state^75^ and overcome replication stress-induced senescence^76^. Specific to mammals, some cell types are consistently polyploid across species while others possess variable amounts of DNA dependent upon the species and the environmental stress^13^. Hepatocytes are polyploid in rodents, large animals and humans^5,7^ and undergo DNA reduction in response to environmental stress to regenerate the liver^77^. Although mechanisms of DNA reduction are not well understood, aneuploidy to increase genetic diversity and resistance to chronic liver injury occurs frequently in human (30-90%) and mice (60%)^60,78^. Whereas aneuploidy can sometimes be maladaptive leading to oncogenesis or cell death, polyploidy generally allows for preservation of normal cellular function^79,80^. Tetraploidy can arise from “mitotic catastrophe” when a proliferating cell fails to undergo cytokinesis following DNA replication, but in general this primarily leads to cell cycle arrest^81^. In rare cases, a select polyploid cell can escape treatments intended to promote senescence such as chemotherapeutic treatment intended to arrest tumor progression^82^. The relevance for polyploidy as a mechanism for avoiding senescence and preserving cell survival in vivo is the subject of ongoing controversy in the cancer literature^75,76^.

In the cardiovascular system, polyploidy is a normal condition of both vascular smooth muscle cells^83^ and cardiomyocytes where an increased frequency of binucleation in rodents is observed compared to increased DNA content within a single nucleus in large mammals and humans^53^. Variation in cardiomyocyte ploidy between species has been documented^34,53^. Higher frequency of diploid cardiomyocytes correlated with better physiologic recovery to myocardial infarction in a survey of multiple murine strains^16^. Diabetic stress also influenced the ploidy level of cardiomyocytes^84^. Although increased numbers of CICs in response to infarction is evident in mammalian models, only one prior publication examined ploidy of the non-myocyte population and concluded that human CICs are diploid^85^, consistent with our finding that human ckit+ CICs are diploid *in situ* and upon isolation and expansion. While polyploidy is a relatively common occurrence in human cardiac muscle and vasculature, human CICs do not readily adopt polyploidy as is the case for rodents.

The importance of the CIC population in response to aging and pathologic injury is well accepted, but only recently have studies delved further into the subtle difference that distinguish subpopulations. This report is the first to our knowledge that delineates ploidy of CICs rather than cardiomyocytes in the heart. Focusing upon the c-kit+ CIC subpopulation allowed for selection of a small subset of the total CIC pool that presumably represents a cell type with progenitor-like characteristics relative to the entire CIC population including multiple committed cell types. While the ultimate role of polyploidy in the CIC population remains to be determined in future studies, the findings in this report establish the benchmark of distinct DNA content differences between rodent and large mammals. We hypothesize an endogenous response within rodent mammals lacking in larger mammals contributes to the blunting of injury along with enhanced capacity for repair^86^. Future studies regarding fresh isolate human lineage-ckit+ CICs may provide additional insights into differences between endogenous human and rodent samples. The challenge of defining functional differences between the 2n versus 4n ploidy states is complicated by the inherent plasticity of ploidy state: attempts to isolate and separately expand diploid versus tetraploid mouse CSCs were unsuccessful because diploid cells underwent binucleation and growth arrest, allowing tetraploid cells persist (data not shown). Although at present we have not performed a direct diploid to tetraploid comparison of CSCs from the same species to determine *in vitro* and *in vivo* adoptive transfer functional activity, our findings provide a compelling rationale for investigations to incorporate polyploidization into cardiac regenerative medicine by induction of human CSC tetraploidization as a method to enhance functional activity.

## Methods

### Mouse CSC and BMSC isolation

All procedures and experiments involving mice were conducted by observing ethical guidelines for animal studies as approved by the SDSU Institutional Animal Care and Use Committee. Mouse CSCs were isolated from both male and female mice for initial karyotype and ploidy observations and maintained as previously described^87^. Cultured male FVB CSCs were used at passage 11 and 100 for experiments unless otherwise stated. Bone marrow stem cells (BMSCs) were isolated from 12-week-old male and female FVB mice by flushing the femur and tibiae with 5% Fetal Bovine Serum in PBS through a 40-µm filter and centrifuged (10 minutes, 600g, 4°C) for initial ploidy analysis with experiments performed on male FVB BMSC for a consistent control against mCSCs. Cells were suspended in 96-well U-bottom plates with media consisting of STEMSPAN, 1% PSG, Flt-3 and SCF: 50ng/ml and IL-3, IL-6 and TPO: 10ng/ml. Media was changed every two days and cells were used at passage 2-4 for experiments unless otherwise stated.

### Human CSC isolation

Samples were received from consenting patients with institutional review board (IRB) approval following NIH guidelines for human subjects’ research. Human CSCs were isolated and maintained as previously described^88^. Karyotype was performed on both male and female CSCs. Cultured hCSCs from three different human tissue samples were used at passage 5 and the respective higher senescence passage (p. 14-24, dependent on sample) for experiments unless otherwise stated.

### Cell karyotype

All cells were G-banded and karyotyped by *Cell Line Genetics* (Madison, WI) except for Sprague Dawley Rat CSCs (karyotyped by Emory University Hospital Oncology Cytogenetics Laboratory, Atlanta, GA) and Feline CSCs (karyotyped by KaryoLogic, North Carolina Biotech Center, Durham, NC). Cells were maintained according to laboratory protocol until prepared for live cell one-day delivery shipping according to an identical preparation protocol by service provider (for example, see: https://www.clgenetics.com/wp/wp-content/uploads/2014/06/Mailing-Live-Cultures-4.17.14.pdf).

### Proliferation assay and cell doubling time

Cell proliferation was determined using the CyQuant Direct Cell Proliferation Assay (Life Technologies, Carlsbad, CA) according the manufacturer’s instruction and as previously described^89^. Population doubling times were calculated using the readings from CyQuant Direct Proliferation Assay and use of a population doubling time online calculator (http://www.doubling-time.com/compute.php).

### Flow cytometry

Flow cytometry to measure DNA content in terms of fluorescence intensity is well documented^90,91^. Suspended cells were fixed using 70% ethanol and kept at −20°C for at least 24 hours prior to measuring DNA content using propidium iodine (PI) with RNAse (BD Biosciences, San Jose, CA). Approximately 50,000 cells were centrifuged at 1300 rpm for 5 minutes at 22°C; the ethanol was removed and the cells were resuspended in 200 μL PI and kept at 37°C in the dark for 10 minutes. 10,000 events per sample were recorded with doublets and unstained events removed from analysis. PI fluorescence intensity was measured using a BD FACSCanto. To sort fresh isolate cells by DNA content, cells are labeled with Vybrant DyeCycle Stain (ThermoFisher, Carlsbad, CA) 1:60,000 in 300μL Wash Buffer (PBS with 0.5% BSA) at 37°C in the dark for 10 minutes. Cells are sorted based on FI of Vybrant dye using a BD FACSAria II, sorting at 9μL/sec. Data is analyzed using FLOWJO (Ashland, OR).

### Immunocytochemistry

Stem cells were placed at a density of 10,000 per well of a two-chamber permanox slide and were fixed using 4% parafamaldihyde for 10 minutes at room temperature. Cells were permeabilized using 0.03M glycine for 5 minutes followed by 0.5% Triton-X 100/PBS for 10 minutes to reduce non-specific binding. Cells were then blocked with TNB for 30 minutes at room temperature followed by primary antibody in TNB labeling overnight at 4°C. The next day, cells were rinsed with PBS then labeled with secondary antibody and/or phalloidin in TNB for 1.5 hours. Subsequent tyramide amplification was performed as necessary. The cells are then rinsed with PBS, DNA is stained with 1 mg/ul 4’,6-diamidino-2-phenylindole (DAPI) (ThermoFisher, Carlsbad, CA) 1/10,000 in PBS for 5 minutes and Vectashield mounting medium is applied (Vector Labs, Burlingame, CA). Slides were visualized using a Leica TCS SP8 confocal microscope. Primary and secondary antibodies used are listed in Supplement Table 2.

### Immunohistochemistry

Mouse hearts were retroperfused, removed from the animal and fixed in 10% formalin overnight. Tissue samples from other organs or heart sections from larger mammals were also formalin fixed overnight. Tissue was then treated with 70% ethanol prior to paraffin embedding using a Leica eASP300 enclosed tissue processor (Leica Biosystems, Buffalo Grove, IL). Paraffin processed tissues were cut into 5 micron section and slide mounted using a HM 355S Automatic Microtome (Thermo Fisher Scientific, Waltham, MA). Heart sections were deparaffinized and incubated with primary and secondary antibodies as previously described^92^. Subsequent tyramide amplification was performed as necessary. The cells are then rinsed with PBS, DNA is stained with 1 mg/ul 4’,6-diamidino-2-phenylindole (DAPI) (ThermoFisher, Carlsbad, CA) 1/10,000 in PBS for 5 minutes and Vectashield mounting medium is applied (Vector Labs, Burlingame, CA). Slides were visualized using a Leica TCS SP8 confocal microscope. Primary and secondary antibodies used are listed in Supplement Table 2.

### In situ confocal microscopy imaging

Measuring fluorescence intensity of the DAPI signal is a known protocol to measure a relative quantity of DNA^93,94^. Briefly, the cells were imaged with a Leica SP8 confocal microscope and a 40X water Objective (Leica, Buffalo Grove, IL), maintained with a 1 Airy unit pinhole size. The DAPI wavelength excitation is 405 and the two hybrid-photomultiplier tube (HyD) emission bandwidth was set at 412-452 nm, at a power intensity of 1.5W and gain of 150V. The CENPB probe is tagged with a Cy3 fluorophore, excited at 561nm, with an emission bandwidth set at 571-620nm under a power intensity of .25W and gain of 60V. The format of the acquisition was 1024X1024 at 400 Hz. The zoom for locating ckit+ cells in tissue was kept at 1, until located. All cells scanned, in both tissue and cell culture were zoomed at a factor of 4, to create a pixel dimension of 0.071μm in both the x and y plane and 0.426 μm in the z-plane during the z-stack scanning. 5 line averages and a single frame average occurred in a bidirectional manner at a frame rate of 0.773/s. Each stack dimension was kept at an optimized system option, with a step size of 0.42 μm, with 19-25 slices included per z-stack. Z-stacks were then processed.

### Cytospun cell cultures

Cultured mCSCs and mBMSCs were suspended at 5 × 10^5 cells/mL in PBS and 200uL was loaded into EZ Cytofunnels (ThermoFisher, Carlsbad, CA) within a Shandon Cytospin 4 (ThermoFisher, Carlsbad, CA). Cells were spun down for 3 minutes on low acceleration and then fixed onto poly-lysine coated slides using Shandon Cell-Fixx fixative (ThermoFisher, Carlsbad, CA). Cells were dried overnight and prepared for immunocytochemistry with phalloidin and DAPI at 1:10,000 for 5 minutes. Confocal imaging utilized a 63x oil objective (Leica, Buffalo Grove, IL), with zoom factor of 0.75, 1.1 Airy unit pinhole, and HyD emission bandwidth of 415-510 at power intensity of 1.5W and gain of 150V. The pixel dimensions were 0.241um in the x and y plane and 0.357um in the z-plane during z-stack acquisition; step size was optimum at 0.35um with 28-30 slices per z-stack.

### Z-stack analysis of DNA content

Z-stacks were reconstructed and analyzed using the Leica SP-8 software (Leica, Buffalo Grove, IL). To begin a region of interest (ROI) was precisely drawn around the widest x-y dimension of each cell nucleus with care taken to not over or under define the ROI. Statistics of the z-stack ROI, the mean fluorescence intensity value of the processed pixels and the ROI area, defined in pixel, are then multiplied to define a pure fluorescence intensity value for the ROI at the given dimension, and consistent gain and intensity. The fluorescence intensity (a unitless measurement) is reported as a relative quantity of DNA content per nucleus. The number of cCICs identified per in situ section varied, with analysis including 6-10 cCICs and 10-25 CMs per experiment. Human tissue was from a single section in the left ventricle while murine tissue included all regions of the heart and cells were selected based on identification of cCICs, which were located throughout the heart.

### Immunoblot sample preparation and experiment

CSCs samples were collected in 1X sodium dodecyl sulfate (SDS) sample buffer with protease and phosphatase inhibitors. Cell lysates were boiled for 10 minutes and stored at −80°C. The analysis for each protein was based on 3 or more separate immunoblots. Within each individual blot, samples were prepared from 3 or more independent cell lysates at low and high passage points. Each sample was normalized to the housekeeping protein GAPDH, and the three samples at low or high passage were averaged. Averaged protein quantities were statistically compared using a t-test analysis based on N=3 separate experimental runs (except for human p53 and S15 phosphorylated p53, which was N=4). Proteins were loaded into 4-12% Bis-Tris gel (Thermo Fisher Scientific, Waltham, MA) and run in 1X MES SDS running buffer (Thermo Fisher Scientific, Waltham, MA) at 150 V for 1.5 hours on an electrophoresis apparatus (Invitrogen, Carlsbad, CA). Separate proteins were transferred to a polyvinylidene difluoride membrane in 1X transfer buffer (Thermo Fisher Scientific, Waltham, MA) then blocked with Odyssey Blocking Buffer (TBS) (LI-COR, Lincoln, NE) for 1 hour. After blocking, membrane was incubated with primary antibodies in Odyssey Blocking Buffer at 4°C overnight. The next day, the membrane was washed with 1X TBST 3 times at 15 minutes, room temperature on an orbital rocker. The membrane was then incubated with secondary antibodies in Odyssey Blocking Buffer for 1.5 hours at room temperature on an orbital rocker followed by three washes of 1X TBST for 15 minutes a wash, room temperature on an orbital rocker. The membrane was then scanned using an Odyssey CLx (LI-COR, Lincoln, NE). Primary and secondary antibodies used are listed in Supplement Table 2.

### mRNA isolation, cDNA synthesis and quantitative PCR

RNA was enriched using the Quick RNA Mini Prep kit from ZymoResearch according to the manufacturer instructions. Reverse transcriptase was performed using protocol for the iScript cDNA Synthesis Kit (BIORAD, Hercules, CA). qRT-PCR was read after incubation of cDNA, primers (100nM) and IQ SYBR Green Supermix (BIORAD). Data was analyzed using the ∆∆C(t). PCR Arrays for a generic cell cycle and p53 signaling analysis were purchased from BioRad (Hercules, CA) All other primer sequences were selected using primer-blast, verified through an oligo calculation, purchased from Thermo Fisher Scientific (Waltham, MA) and listed in Supplement Table 3.

### scRNAseq cell preparation and sequencing

Preparation of cells for single cell gene expression was performed according to manufacturer specification (10X Genomics, Pleasanton, CA). To optimize gem tagging efficiency, 2000 cells were loaded per group into the Chromium system and indexed sequencing libraries were constructed according to the manufacturer protocol using the Chromium Sing Cell 3’ Library & Gel Bead Kit v2 (10X Genomics, Pleasanton, CA). Each library underwent quality control screening using a Bioanalyzer (Agilent Genomics, Santa Clara, CA) and quantified by quantitative PCR (KAPA Biosystems Library Quantification Kit for Illumina platforms) and Qubit 3.0 with dsDNA HS Assay Kit (Thermo Fisher Scientific, Watham, MA). Sequencing libraries were loaded at 2pM on a Illumina HiSeq2500 with 2X75 paired-end kits using the following read length: 98 bp Read 1, 8 bp i7 Index and 26 bp Read 2.

### scRNAseq analysis

The raw data was processed with the Cell Ranger pipeline (10X Genomics; version 2.0). Sequencing reads were aligned to the mouse genome mm10. Preparations derived from two mouse hearts per sample were used to produce 1036 diploid and 628 tetraploid freshly isolated cells for analysis. Cells with fewer than 1,000 genes or more than 10% of mitochondrial gene UMI count were filtered out and genes detected fewer than in three cells were filtered out^95^. Altogether, 1,664 cells and 15,579 genes were kept for downstream analysis using Seurat R Package (v2.3.4). Approximately 2,027 variable genes were selected based on their expression and dispersion. The first 12 principal components were used for the t-SNE projection^96^ and unsupervised clustering^95^. Gene expression pathway analysis was performed using clusterProfiler^97^. scRNA-Seq data generated was uploaded to the Gene Expression Omnibus (GEO submission GSE122057, released November 2, 2018).

### Myocardial infarction

Twelve-week old female FVB were sedated and myocardial infarction was produced by ligating the left anterior descending (LAD) branch of the coronary artery using a 8-0 suture (Ethicon, Somerville, NJ). Echocardiography was performed under mild isolflurane sedation (0.5%-1.5%) using a Vevo 2100 Linear Array Imaging ultrasound (FUJIFILM Visual Sonics, Toronto, ON, Canada). Cardiac function was analyzed is the parasternal long axis view by tracking the endocardium the supplied analysis software to obtain end systolic volume, end diastolic volume, ejection fraction and heart rate. Noninvasive assessment of cardiac function in the parasternal long axis view was performed at baseline, the day before surgery, and the day before harvesting to verify similar infarction size and cardiac function.

### Statistical analyses

All data are expressed as mean +/− standard error of mean. Statistical analyses were performed within and between group comparisons, student t-test, one- and two-way ANOVA was applied with Bonferroni post-hoc test, when applicable. using Graph Pad Prism v5.0 (GraphPad Software, La Jolla, CA). A value of less than 0.05 was considered statistically significant.

## Acknowledgments

We thank Cameron Smurthwaite of the San Diego State University Flow Cytometry facility for flow cytometry assistance with sorting fresh cCIC isolates. We thank Dr. J. Hare (University of Miami) for providing Yorkshire swine CSC samples as well as Gottigen swine cardiac tissue samples. We also thank Dr. Marcel Rota (New York Medical College) for his support by providing Canine cardiac tissue samples. Adult normal human tissue samples were provided by the NCI Cooperative Human Tissue Network (CHTN). Lastly, we thank the staff at Sharp Hospital, in particular Chris Kohlmeyer and Donna Small, for bridging the collaborative arrangement between Sharp Hospital and the Sussman Laboratory. K.M.Broughton is supported by NIH grant F32HL136196. M.A. Sussman is supported by NIH grants: R01HL067245, R37HL091102, R01HL105759, R01HL113647, R01HL117163, P01HL085577, and R01HL122525, as well as an award from the Fondation Leducq.

## Author Contributions

KMB and MAS conceived the project. KMB SM, PQ, BJW, DK, MM and TK performed CSC and BMSC isolations. KMB, SM, PQ and MD prepared samples for karyotype analysis. KMB prepared samples and conducted CANTO flow cytometry, western blot, and qPCR experiments. KMB, TK, NN, MR and NG prepared immunocytochemistry and immunohistochemistry samples and conducted analysis. KMB, OE and TYK prepared samples and performed analysis for scRNAseq. KMB and MAS analyzed data and wrote the manuscript. All authors provided critical feedback on the manuscript.

