## Supplement for "Cardiac interstitial tetraploid cells that escape replicative senescence are found in rodents but not large mammals"

**Supplement Table 1**

| | Average Expression (TPM) | | | | |
|---|---|---|---|---|---|
| Gene | 2N | 4N | Cluster | Cluster FC | Adjusted p.val |
| Col3a1 | 8.153936 | 3.169273 | 2N | 6.115647 | 5.69E-40 |
| Mmp2 | 4.410567 | 2.036908 | 2N | 3.780194 | 8.43E-37 |
| Dcn | 118.3096 | 86.85434 | 2N | 2.023212 | 1.97E-18 |
| Thbs1 | 5.091703 | 1.432918 | 2N | 8.276304 | 3.15E-40 |
| Pecam1 | 1.575726 | 3.378566 | 4N | 3.393048 | 4.42E-22 |
| Cdh5 | 2.149346 | 4.720082 | 4N | 3.95174 | 5.39E-24 |
| Cd36 | 5.877553 | 13.63402 | 4N | 5.689655 | 6.00E-32 |
| Tcf15 | 2.053154 | 3.945359 | 4N | 3.035808 | 1.08E-16 |
| Gpihbp1 | 5.113292 | 10.25305 | 4N | 4.07543 | 7.87E-21 |

**Method Tables**

| Use: Immuno-histochemistry | Company | Antibody Dilution |
|---|---|---|
| Goat anti-CD117 (ckit) | R&D Systems | 1:100 |
| Bovine anti-goat HRP* | ThermoFisher | 1:200 |
| Mouse anti-cardiac troponin T | ThermoFisher | 1:100 |
| Mouse Tryptase | Abcam | 1:100 |
| Rabbit Desmin | Abcam | 1:100 |
| Donkey anti-mouse 647 | ThermoFisher | 1:200 |
| Donkey anti-mouse 488 | ThermoFisher | 1:200 |
| Donkey anti-Rabbit 647 | ThermoFisher | 1:200 |
| Donkey anti-Rabbit 488 | ThermoFisher | 1:200 |
| Tyramide TRITC | ThermoFisher | 1:100 |
| DAPI (1mg/uL) | Sigma-Aldrich | 1:10000 |

| Use: Immuno-cytochemistry | Company | Antibody Dilution |
|---|---|---|
| Goat anti-CD117 (ckit) | R&D Systems | 1:200 |
| Bovine anti-goat HRP* | ThermoFisher | 1:400 |
| Phalloidin 488 | ThermoFisher | 1:400 |
| Tyramide TRITC | ThermoFisher | 1:100 |
| DAPI (1mg/uL) | Sigma-Aldrich | 1:10000 |
| Topro3 | ThermoFisher | 1:10000 |

| Use: Western Blot | Company | Antibody Dilution | Molecular Weight |
|---|---|---|---|
| Chicken GAPDH | Abcam | 1:500 | 37 kDa |
| Rabbit Cdc25c | Abcam | 1:500 | 53 kDa |
| Rabbit P53 | Abcam | 1:500 | 53 kDa |
| Rabbit P53 Phospho-S15 | Abcam | 1:300 | 53 kDa |
| Rabbit MDM2 | Abcam | 1:300 | 75, 90 kDa |
| Rabbit TRIM13 | ProteinTech | 1:500 | 47 kDa |
| Rabbit TRIM25 | Abcam | 1:500 | 71 kDa |
| Rabbit TRIM28 | ProteinTech | 1:500 | 100 kDa |
| Rabbit TRIM29 | Abcam | 1:500 | 66 kDa |
| Donkey Anti Chicken 700 | Licor | 1:1000 | |
| Donkey Anti Rabbit 800 | Licor | 1:1000 | |

**Table 2: Antibody List**

| mRNA Primer Gene | Forward 5' - 3' | Reverse 5' - 3' |
|---|---|---|
| Human | | |
| Trim 2 | TTTGCAGGTCCCCATTTTGC | GCCATAGAGTGGGTCAGCAG |
| Trim 13 | CACTAGCCGGAGTAGCCTCT | TCCACAAGGAATTCCGCACA |
| Trim 19 | ACAACGACAGCCCAGAAGAG | CGAGCTGCTGATCACCACAA |
| Trim 25 | TACATCCCCGAGGTGGAACT | GGAGACCTTCTTCACAGGGC |
| Trim 28 | TCTTGGGCTCTGGAGAGTGA | TTGGTCCAGGCATTGAGGTC |
| Trim 29 | TTTCCCTCCTGCTCTTGCTG | AAGTTCTGCTCCAGGATGGC |
| Mouse | | |
| Trim 2 | AGTTTGCAGGTCCCCACTTT | CCCCAGTCAGCCACAATGAT |
| Trim 13 | CAGTTGGCTGGTGGAGTGTT | CAACACTCGGGGGTCATCAA |
| Trim 19 | AGCTGCTCACCAGAGGTTTC | AAGCCTCCTGCTCAAGGTC |
| Trim 25 | AAGCAACTTCCCCTGATGCC | TTGTTGTGCCAGGCAGAGAT |
| Trim 28 | GGTGAGAAGCGTCCGGC | GGTTCAGAGCACTCCACACA |
| Trim 29 | AAAGGCTTTCCCTCCCTCCT | CAGAGACTGTGTGAGGGCAG |

**Table 3: qRT-PCR Primer List**

**Supplement Figure Titles and Legends**

**Supplement Figure 1: Mononuclear tetraploid content unique to rodent cardiac stem cells is verified with multiple karyotypes.**
G-band karyotype analysis was performed on cultured CSCs to reveal tetraploid content of rodent C57 mouse (A) and CAST mouse (B) with diploid content of feline (C), fetal human (D,E) and adult LVAD patient (F) CSC samples.

**Supplement Figure 2: Morphology and proliferation rate of CSCs is similar within species.**
Immunocytochemistry of CSC verify mononuclear content from multiple human samples (A-C), swine samples (D-F), rat samples (G-L) and mouse samples (M-O) with zoomed in images (A'-O'; scalebar = 100um). Surface area increased in human CSCs from an LVAD patient compared to normal controls (P), while surface area was consistently smaller for CSCs isolated from swine (Q), rat (R), and mouse (S) samples. CSCs from all samples demonstrated an overall spindle appearance based on major to minor axis ratio. Proliferation rates were consistent within each animal species (T-W).

**Supplement Figure 3: Tetraploid CSCs proliferate faster than diploid CSCs.**
Proliferation rates were compiled within human and mouse species at early and late passage points, demonstrating mCSCs proliferate at a similar rate to hCSCs between days 0 and 3 but is statistically increased by day 4 (A). Correlated with proliferation rate, doubling time between day 0 to day 4 after plating, demonstrated mCSCs proliferate faster, but not statistically significant, than hCSC at early passage; last passage hCSCs demonstrate replicative senescence (B). ***$P<0.001$. Data are presented as Mean±SEM and analyzed using a two-way ANOVA with Bonferroni post-hoc test (A,B).

**Supplement Figure 4: Tetraploid murine CSC content remains stable and CSCs proliferate faster over increased passages**

Immunocytochemistry image of FVB CSCs at low (A) and high (B) passage demonstrate mononuclear content (scalebar = 200um). Surface area of FVB CSCs at low and high passage is comparable (C). Flow cytometry of DNA content stained with propidium iodine from FVB CSCs at passage 12,25,50 and 100 demonstrates consistent and stable tetraploid content (D). Proliferation of FVB CSCs at passage 10, 50 and 100 demonstrate a correlative increase with passage point (E), and faster doubling times (F). Cell cycle transcription expression is statically significantly increased in Ccna2, Ccnb2 and Cdc25c in high passage compared to low passage FVB CSCs, statistically analyzed using t-test per gene (G). Protein expression of Cdc25c increases with passaging and verifies cell cycle gene expression and increased proliferation rates (H). *P<0.05; **P<0.01; ***P<0.001. Data are presented as Mean±SEM and analyzed using a t-test between each gene or protein (C,G H), or one-way (F) or two-way (E) ANOVA with Bonferroni post-hoc test.

**Supplement Figure 5: Murine CSCs downregulate the p53 pathway with increased passages.**

P53 pathway associated transcription markers and proteins were analyzed in early passage compared to late passage for human and murine CSCs. High passage, compared to low passage, human CSCs display increased transcription for apoptosis marker BCL2 and endothelial growth factor receptor KDR, both found in the p53 pathway (A). Likewise, high passage human CSCs display unchanged p53 protein levels but decreased phosphorylated S15 of p53 and MDM2 protein (B,C). In high passage FVB murine CSCs, transcription gene Hspb1 for making proteins that block programmed cell death and Cdk2, to regulate the cell cycle from S to G2, are upregulated while senescence transcription genes for p53 (Trp53), p16 (Cdkn2a), and p21 (Cdkn1a) are down regulated (D). Protein levels of p53, phosphorylated S15 p53 and MDM2 are also down in high passage FVB CSCs (E,F), verifying transcriptional results. *P<0.05; **P<0.01; ***P<0.001. Data are presented as Mean±SEM and analyzed using a t-test between each gene or protein (A-F).

**Supplement Figure 6: Murine CSCs increase negative p53 feedback loop with increased passages.**

TRIM family transcription markers involved with potentiation of p53 (TRIM 13,19) or inhibition (TRIM 25,28,29) via the Mdm2-p300-p53 complex were analyzed at the transcriptional and protein levels at early and late passage human and murine CSCs. Transcription for TRIM 13 was upregulated and TRIM 25 and 28 were downregulated in high, compared to low, passage human CSCs (A). Transcription for TRIM 13 and 19 was significantly down, while TRIM 25, 28 and 29 were significantly up in high, compared to low, passage FVB CSCs (B) Protein for TRIM 13 (C) was slightly higher, while TRIM 25 and 28 (D) was significantly lower in high, compared to low, passage human CSCs. Protein for TRIM 13 was significantly lower (E), while TRIM 28 and 29 was slightly lower in high, compared to low, passage murine CSCs (F). *P<0.05; **P<0.01; ***P<0.001. Data are presented as Mean±SEM and analyzed using a t-test between each gene or protein (A-F).

**Supplement Figure 7: Fresh isolate murine diploid Lin-Ckit+ CICs primarily represent fibroblast transcriptional profiles.**

Single cell RNA sequencing was used to identify cellular profiles of the freshly isolated lin-ckit+ CICs from adult FVB mice sorted based on diploid and tetraploid state. Results demonstrate the diploid population has significantly more cells positive for fibroblast-related markers based on Col3a1, Mmp2, Dcn and Thbs1, as shown with tSNE (A), percentage of cells expressing each gene (B), and violin plots showing heterogeneity for gene expression (C).

**Supplement Figure 8: Fresh isolate murine tetraploid Lin-Ckit+ CICs primarily represent endothelial transcriptional profiles.**

Single cell RNA sequencing was used to identify cellular profiles of the freshly isolated lin-ckit+ CICs from adult FVB mice sorted based on diploid and tetraploid state. Results demonstrate the tetraploid population has significantly more cells positive for endothelial-related markers based on Pecam1, Cdh5, Cd36, Tcf15, and Gpihbp1, as shown with tSNE (A), percentage of cells expressing each gene (B), and violin plots showing heterogeneity for gene expression (C).

**Supplement Figure 9: *In situ* immunohistochemistry DAPI concentration selection.**

Adult C57 mouse heart sections were prepared for immunohistochemistry and stained for DAPI (1mg/ul) at dilution of 1:1000 or 1:10,000 with a time course of 5, 15, 30 minutes and 1, 4, 16 hours. Topro3 was stained 1:10,000 and used as a second control to stain DNA. Z-stacks were created and analysis focused on cardiomyocytes. Analysis demonstrated more variability of fluorescence intensity for DAPI with 1:1000 dilution at all time points, while 1:10000 dilution demonstrated consistency for the 5-15 minute time course and increased variability with longer staining time (A). Visual representation confirmed DAPI dilution at 1:1000 yielded non-specific fluorescence outside of the tissue section (B), while DAPI dilution at 1:10,000 stained within the tissue, without non-specific fluorescence outside of the tissue, and aligned with Topro3 stain (C). Fluorescence intensity of DNA content in cCICs and CMs from human cardiac fetal and normal adult tissue, stained using a 1:10,000 DAPI concentration for 5 minutes (D).

### Supplement Figure 1

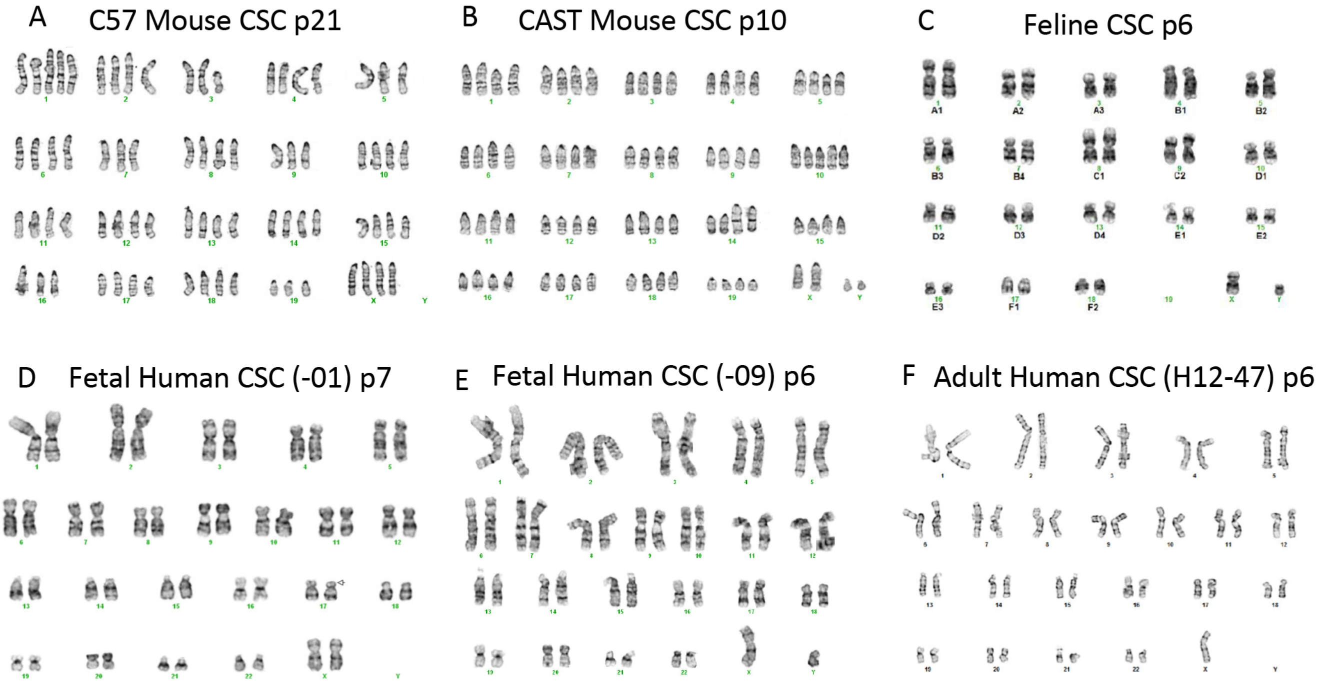

### Supplement Figure 2.1

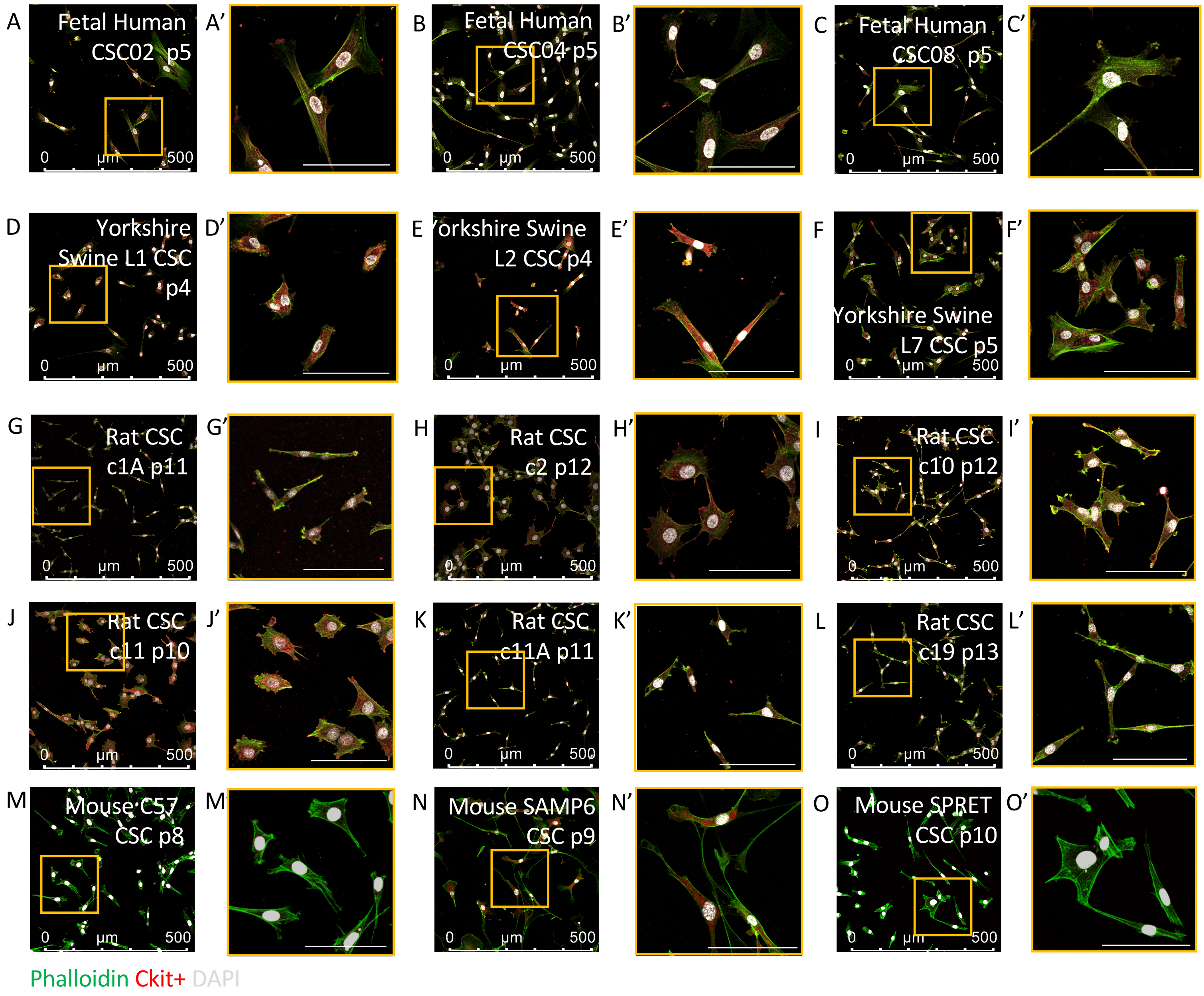

### Supplement Figure 2.2

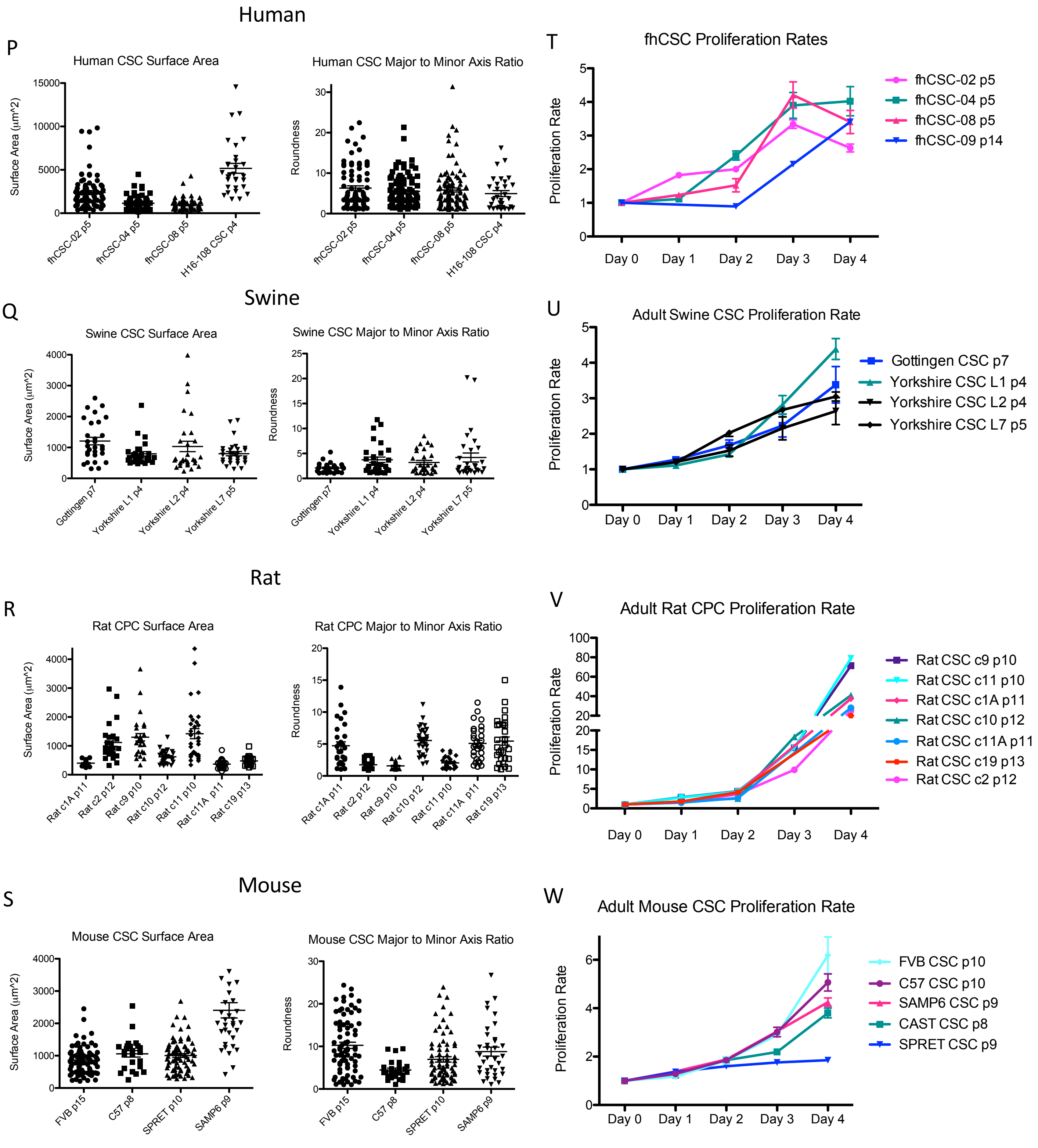

### Supplement Figure 3

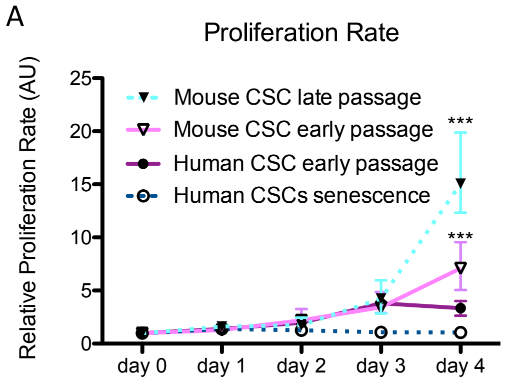


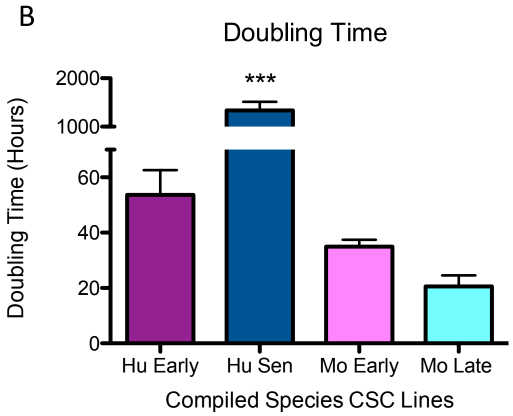

### Supplement Figure 4

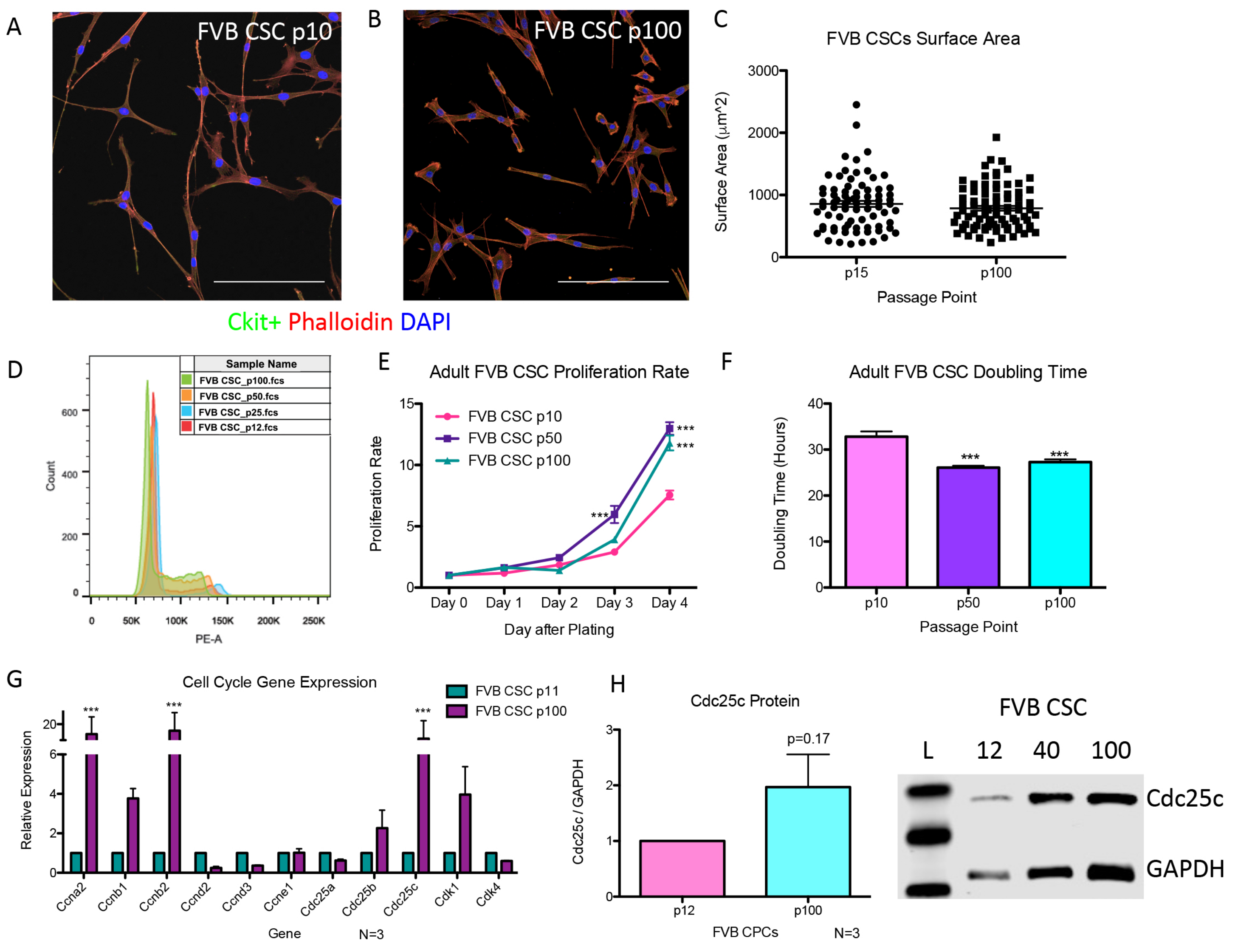

### Supplement Figure 5

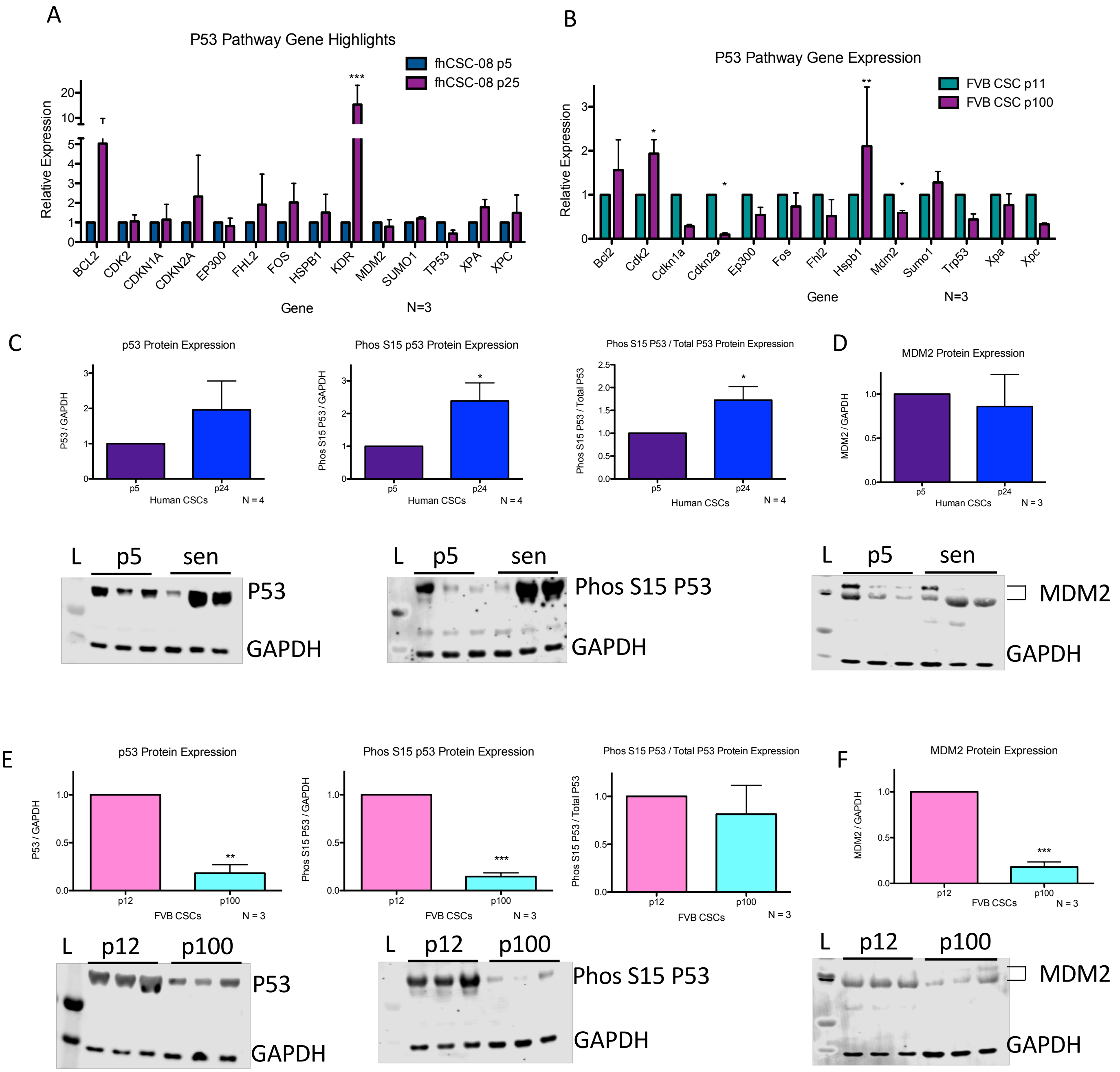

### Supplement Figure 6

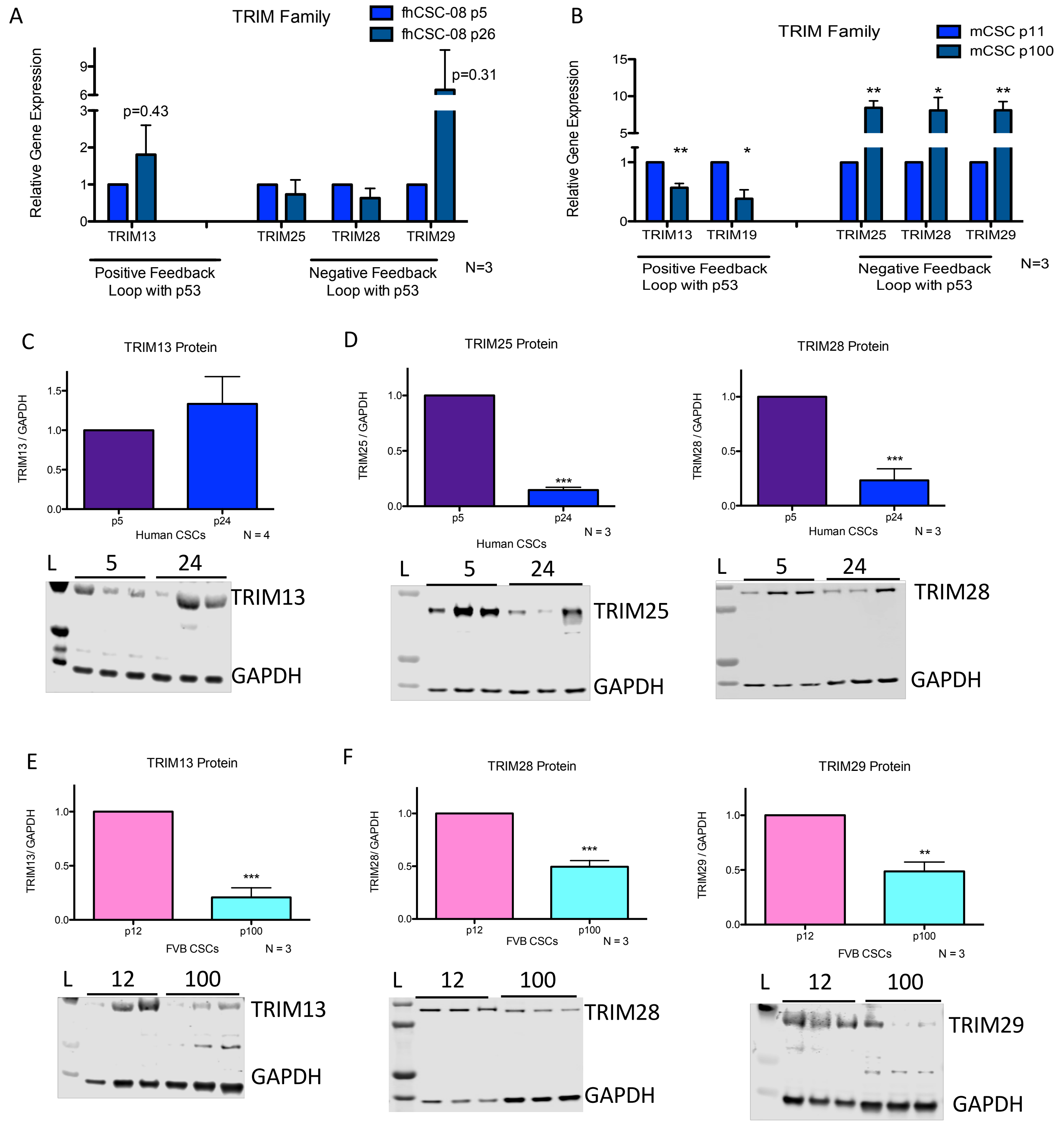

#### Supplement Figure 7

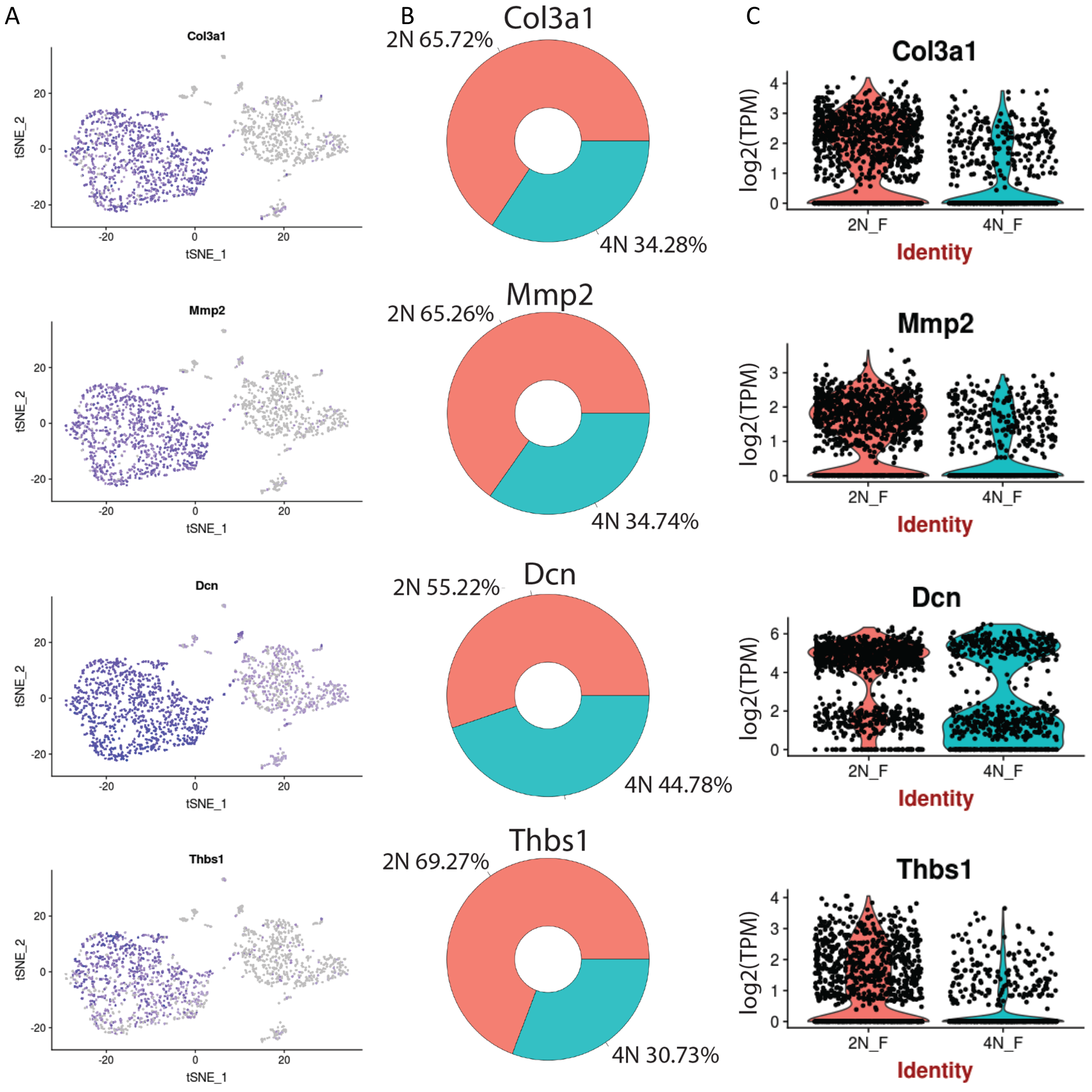

#### Supplement Figure 8

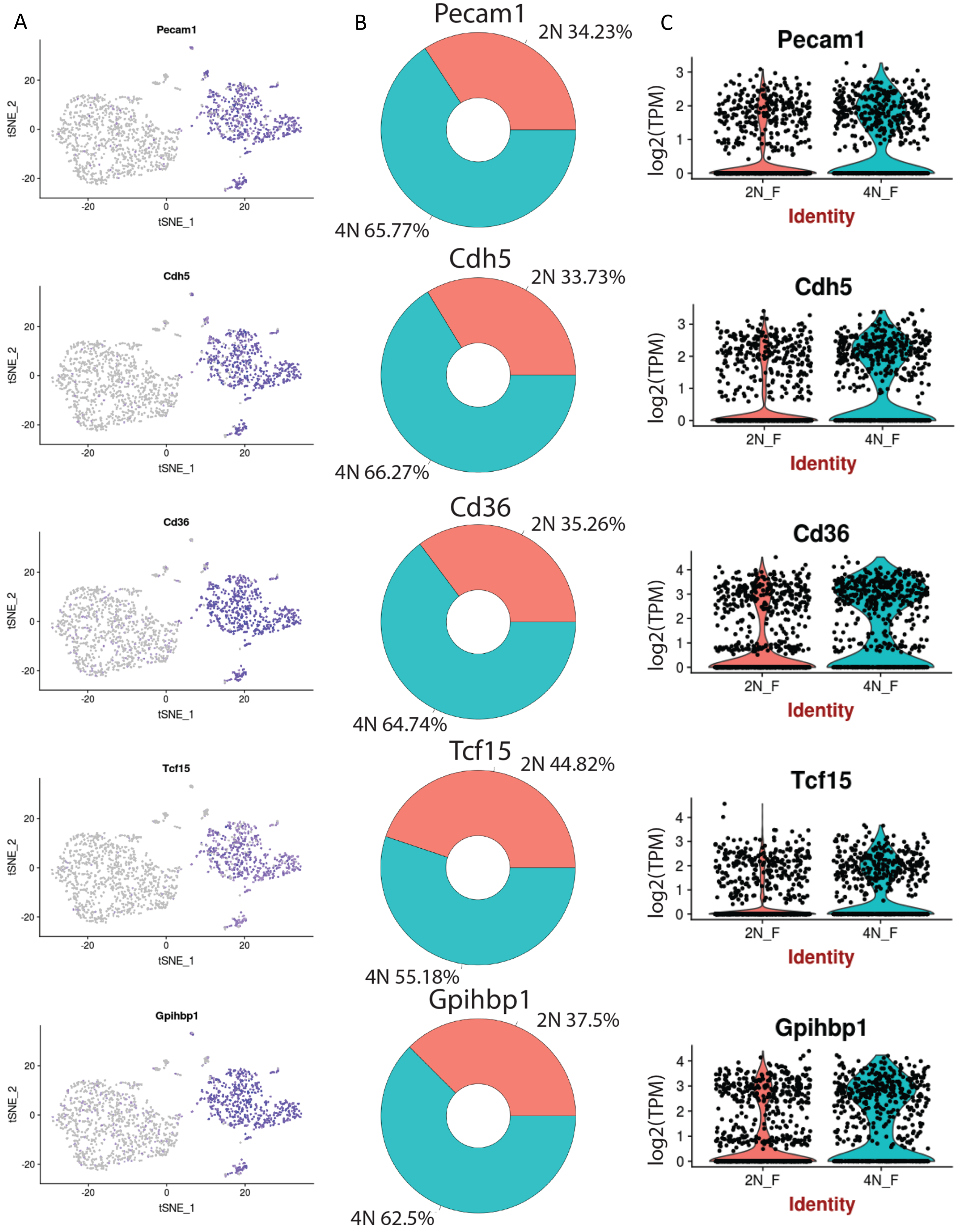

### Supplement Figure 9

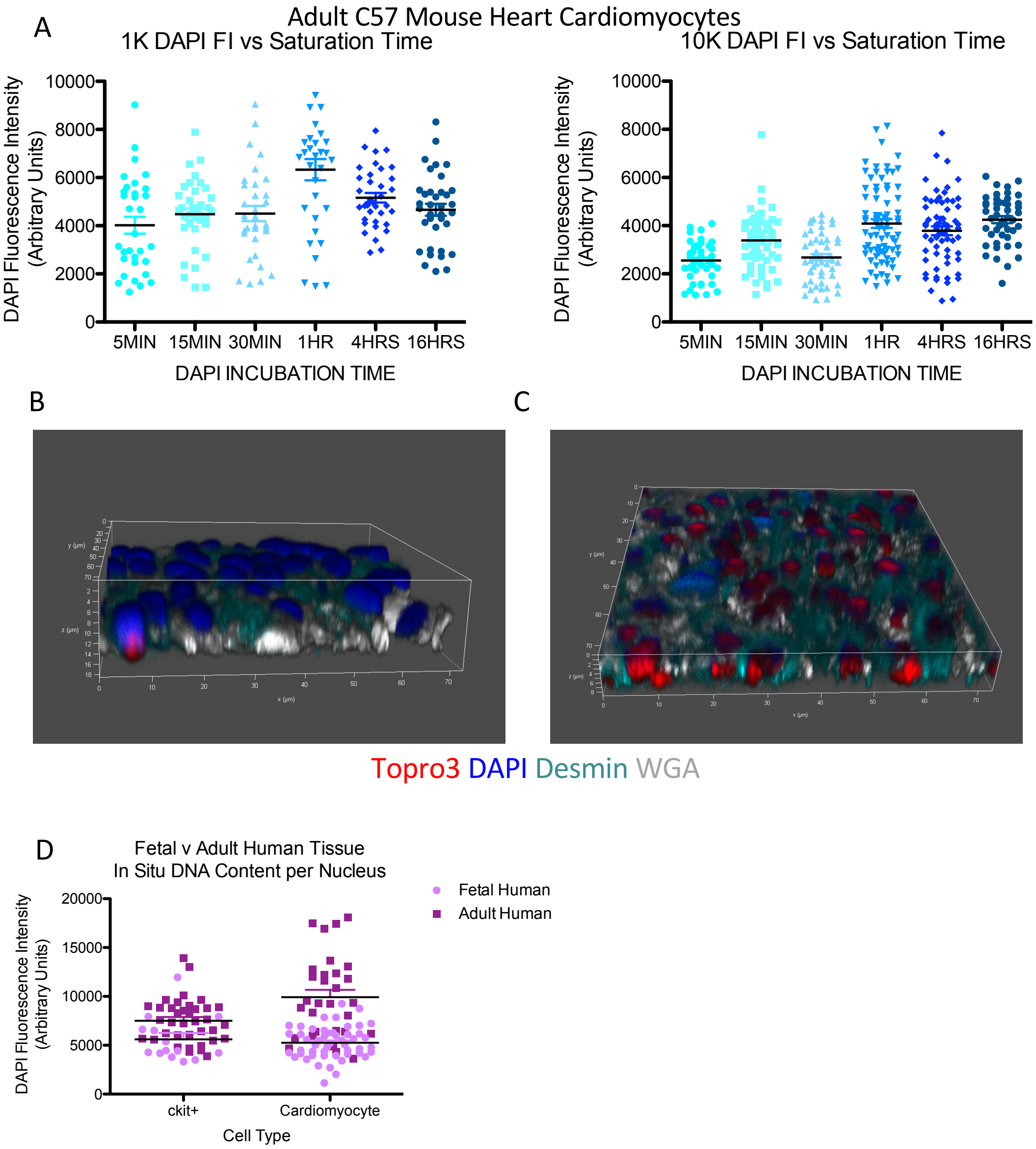
